# Phosphorylation of HP1/Swi6 relieves competition with Suv39/Clr4 on nucleosomes and enables H3K9 trimethyl spreading

**DOI:** 10.1101/2024.10.25.620326

**Authors:** Dana R Kennedy, Joël Lemière, Ahmed AA Amine, Eric W Martin, Catherine Tan, Eric Simental, Julian Braxton, Robert A Maxwell, Bassem Al-Sady

## Abstract

Heterochromatin formation in Schizosaccharomyces pombe requires the spreading of histone 3 (H3) Lysine 9 (K9) methylation (me) from nucleation centers by the H3K9 methylase, Suv39/Clr4, and the reader protein, HP1/Swi6. To accomplish this, Suv39/Clr4 and HP1/Swi6 have to associate with nucleosomes both nonspecifically, binding DNA and octamer surfaces and specifically, via recognition of methylated H3K9 by their respective chromodomains. However, how both proteins avoid competition for the same nucleosomes in this process is unclear. Here, we show that phosphorylation tunes oligomerization and the nucleosome affinity of HP1/Swi6 such that it preferentially partitions onto Suv39/Clr4’s trimethyl product rather than its unmethylated substrates. Preferential partitioning enables efficient conversion from di-to trimethylation on nucleosomes *in vitro* and H3K9me3 spreading *in vivo*. Together, our data suggests that phosphorylation of HP1/Swi6 creates a regime that increases oligomerization and relieves competition with the “read-write” mechanism of Suv39/Clr4, together promoting for productive heterochromatin spreading.

---

Heterochromatin is a gene-repressive nuclear structure conserved across eukaryotic genomes (Grewal 2023). Gene-repression is signaled by the deposition of characteristic histone marks, such as Histone 3 lysine 9 methylation (H3K9me), by enzymes termed “writers”, for example, homologs of the suppressor of variegation 3-9 methyltransferase (Elgin and Reuter 2013) (Suv39, Clr4 in *S. pombe*). Heterochromatin assembly occurs in two steps: Initially, a domain is specified in a process called nucleation. This occurs by tethering of the “writer” directly to the underlying DNA sequence via DNA-bound factors (Jia et al. 2004) or indirectly via RNA intermediates emanating in cis (Bayne et al. 2010; Khanduja et al. 2024; Allshire and Madhani 2018). The subsequent step involves the lateral spreading of heterochromatic marks over varying distances to define a silenced domain (Hamali et al. 2023). This process, once outside nucleation sites, is no longer mediated by sequence-mediated “writer” recruitment and instead requires two essential features: 1. A positive feedback relationship in which the “writer” also contains a specialized histone-methyl binding chromodomain (CD) that recognizes its own product, H3K9me (Zhang et al. 2008; Muller et al. 2016). 2. a “reader” protein (Elgin and Reuter 2013; Noma et al. 2004; Hall et al. 2002), Heterochromatin Protein 1 (HP1, Swi6 in *S. pombe*), that also recognizes H3K9me2/3 via a CD (Jacobs and Khorasanizadeh 2002).

How do HP1 proteins execute their essential function in heterochromatin spreading? One way in which they do so is by directly recruiting the Suv39 methyltransferase to propagate H3K9 methylation (Haldar et al. 2011; Jenuwein and Allis 2001; Aagaard et al. 2000). Second, HP1 proteins oligomerize on H3K9me-marked chromatin, which has been invoked as a mechanism that supports spreading (Canzio et al. 2011). HP1 oligomerization also underlies its ability to undergo Liquid-Liquid Phase Separation (LLPS) *in vitro*on its own or with chromatin (Larson et al. 2017; Sanulli and J Narlikar 2020; Keenen et al. 2021), and condensate formation *in vivo* (Larson et al. 2017; Strom et al. 2017; Sanulli and J Narlikar 2020; Sanulli et al. 2019). This condensate formation may promote spreading by providing a specialized nuclear environment that concentrates HP1 and its effectors (Holla et al. 2020) and/or excludes antagonists of heterochromatin (Larson et al. 2017). The silencing of heterochromatin by HP1 may be coupled to spreading by oligomerization, which likely promotes chromatin compaction and blocks RNA polymerase access (Fischer et al. 2009; Verschure et al. 2005). Silencing may also require oligomerization-independent mechanisms like HP1’s ability to bind RNA transcripts and recruit RNA turnover machinery (Keller et al. 2012; Motamedi et al. 2008).

However, these proposed mechanisms for HP1’s role in spreading do not contend with a central problem, which is that HP1 and Suv39/Clr4 directly compete for the same substrate on multiple levels. This competition can be specific, as HP1 and Suv39/Clr4 have CDs that recognize the H3K9me2/3 chromatin mark (Canzio et al. 2011; Al-Sady et al. 2013). It is also non-specific, as both HP1 and Suv39/Clr4 bind DNA and histone octamer surfaces of the nucleosome substrate (Muller et al. 2016; Al-Sady et al. 2013; Akoury et al. 2019; Canzio et al. 2013; Sanulli et al. 2019; Shirai et al. 2017). How can HP1 promote H3K9 methylation spreading by Suv39/Clr4, but not get in its way? One explanation for managing the specific competition is an observed difference in methylation state preference. Clr4, for example, is more selective for the terminal trimethylated (H3K9me3) state than Swi6 or the other HP1 paralog in *S. pombe*, Chp2 (Al-Sady et al. 2013). However, how the significant H3K9me3-independent nucleosome affinity of Clr4 and Swi6 is coordinated to avoid competition is not clear.

One possible way to regulate competition in spreading is through post-translational modifications of HP1. For example, HP1a, HP1α, and Swi6 are phosphorylated by CKII protein kinases (Shimada et al. 2009; Eissenberg et al. 1994; LeRoy et al. 2009). Phosphorylation of HP1 across species has been shown to regulate multiple of its biochemical activities including LLPS (Larson et al. 2017), specificity for H3K9me (Larson et al. 2017; Nishibuchi et al. 2014), and affinity for nucleic acids (Nishibuchi et al. 2014). In *S. pombe*, several Swi6 *in vivo* phosphorylation sites have been documented in the N-terminal extension (NTE), the CD, and the hinge domain (Shimada et al. 2009), which, when mutated, disrupt transcriptional gene silencing (Shimada et al. 2009). While phosphorylation of HP1 has been recognized as an important regulatory modification for 20 years (Zhao et al. 2001), the mechanisms by which phosphorylation-induced biochemical changes in HP1 direct its cellular activity and coordination with H3K9 “writers” remain unclear.

In this study, we focused on previously identified Swi6 phosphorylation target sites (Shimada et al. 2009) and found that two sites in particular, S18 and S24, are required for the spreading, but not nucleation, of heterochromatin. Spreading defects in Swi6 S18/24A mutants arise from the inability to convert H3K9me2 to H3K9me3 outside creation sites. We show biochemically that the primary role of phosphorylation is to increase Swi6’s oligomerization capacity and, in turn, lower Swi6’s overall chromatin affinity. This lowered affinity preferentially partitions Swi6 onto H3K9me3 nucleosomes, rather than unmethylated nucleosomes, in vitro, and directs Swi6 molecules into nuclear foci and heterochromatin nucleation sites, rather than euchromatic loci and/or the nucleoplasm, *in vivo*. It may appear counter-intuitive that lowered affinity should have this effect. However, since phosphorylation also increases Swi6’s propensity to oligomerize, this ultimately reduces the Swi6 pool available to bind unmethylated sites. We propose that phosphorylation of Swi6 frees up Clr4’s substrates for efficient trimethylation, and thus, spreading.

## Results

### Serines 18 and 24 are necessary for heterochromatin spreading but not nucleation

Previously, several phosphoserines in Swi6 have been shown to play a role in heterochromatin gene silencing (Shimada et al. 2009) (Figure 1A). To address whether the phosphorylation targets play a role in nucleation and/or spreading of heterochromatin, we used our mating type locus (MAT) heterochromatin spreading sensor (HSS) (Figure 1B). The HSS is based on a series of transcriptionally encoded, fluorescent reporters driven by the ade6 promoter, which is sensitive to heterochromatin formation (Greenstein et al. 2022, 2018). We place a SF-GFP reporter into nucleation sites (‘green’), a mKO2 reporter to monitor distal spreading (‘orange’) and a E2C reporter (‘red’) at a euchromatic locus to control for cell-to-cell transcriptional noise (Al-Sady et al. 2016). The HSS thus allows us to separate nucleation and spreading events at single-cell resolution. Specifically, we used a MAT locus HSS with only the cenH nucleator intact (MAT ΔREIII HSS (Greenstein et al. 2018)), which enables us to isolate spreading from one nucleator.

**Figure 1:**
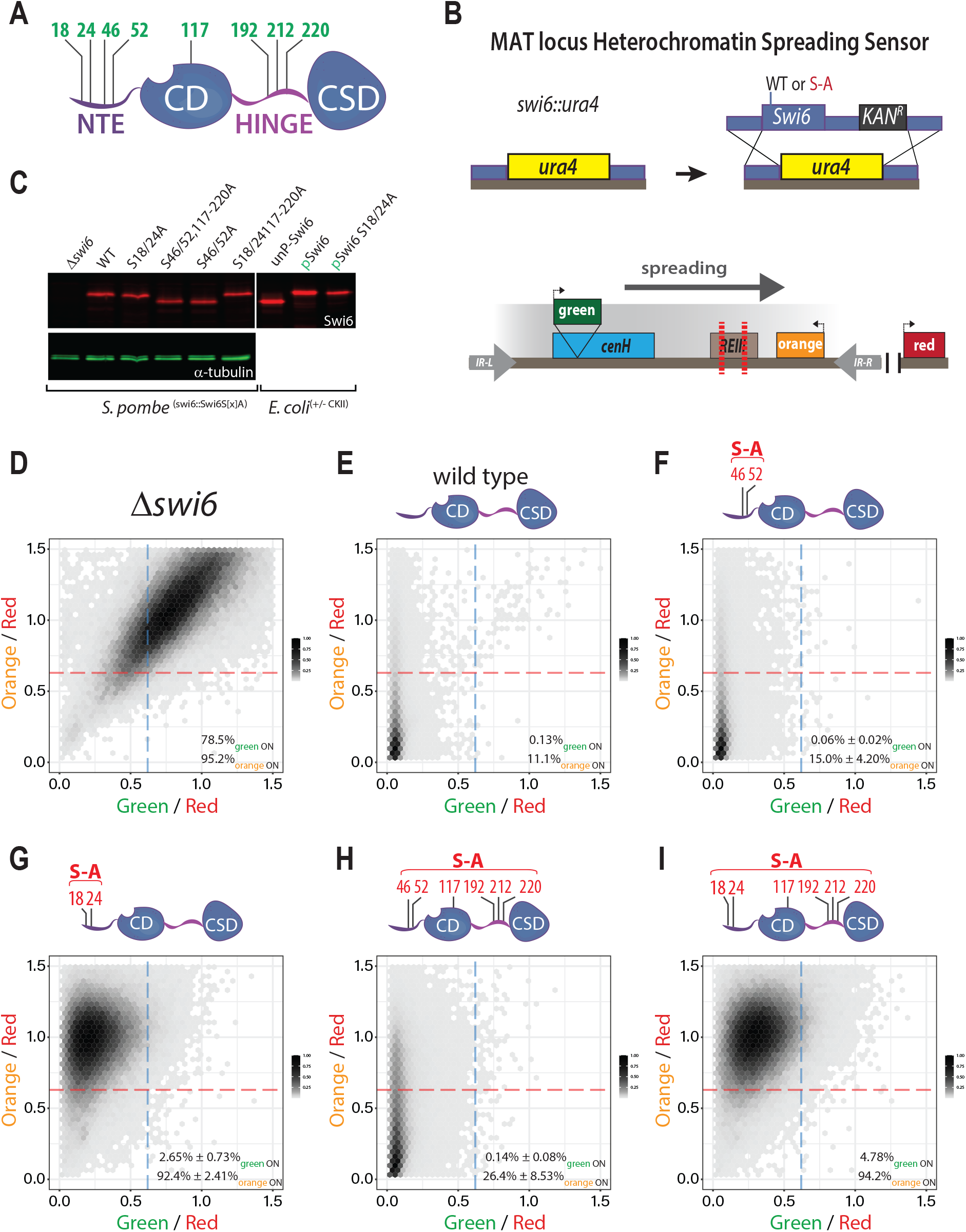
S18 and S24 in Swi6 are required for spreading, but not nucleation of heterochromatin silencing. **A**. Overview of the Swi6 protein domain architecture and previously identified (Shimada *et al*.) *in vivo* phosphorylation sites (green residue numbers). NTE: N-terminal extension; CD: chromodomain (H3K9me binding); HINGE: unstructured hinge region; CSD: chromo-shadow domain (dimerization and effector recruitment). **B**. Strategy for production of *swi6* S-A mutants in the MAT locus *ΔREIII* Heterochromatin Spreading Sensor (HSS, lower diagram) reporter background. **C**. Swi6 levels are not affected by serine (S) to alanine (A) S-A mutations. Total extracts of *swi6* wildtype or indicated mutants were probed with an anti-Swi6 polyclonal antibody, or anti-*α*-tubulin monoclonal antibody as loading control. *In vitro* purified Swi6 that was either phosphorylated (pSwi6) or not (unpSwi6) are run as size controls. Note, not all mutant Swi6 proteins display a band shift even if they retain phosphosites. **D.-I**. 2-D Density hexbin plots examining silencing at nucleation ‘green’ and spreading ‘orange’ reporter in *Δswi6*, wildtype, and indicated S-A mutants. The dashed orange and blue lines indicate the threshold for full expression of the ‘orange’ and ‘green’ reporters, respectively. Indicated percentages and SD represent the fraction of cells above the line.

To query swi6 serine-to-alanine (S-A) mutants in this background, we first replaced the swi6 open reading frame with the ura4 gene (swi6::ura4). Using homologous recombination, we then replaced the ura4 cassette with either wildtype or S-A mutant swi6 open reading frames followed by a kanamycin resistance marker (Figure 1B). We based our S-A mutations on the phosphoserines previously identified in Shimada et al., which include S18, S24, S46, and S52 in the NTE, S117 in the CD, and S192, S212, and S220 in the hinge (Figure 1A). Here, we constructed the following S-A mutants: S18A and S24A (*swi6*^S18/24A^); S46A and S52A (*swi6*^S46/52A^); S46A, S52A, S117A, S192A, S212A, and S220A, (*swi6*^S46/52/117-220A^, “S18/S24 available” ); and S18A, S24A, S117A, S192A, S212A, and S220A (*swi6*^S18/24/117-220A^, from here on *swi6*^6S/A^). These mutants are expressed at similar levels compared to wildtype as assessed by western blot, using a polyclonal anti-Swi6 antibody (Figure 1C, further validated by cytometry in SFigure 6C). Note that not all phospho-site mutants yield an observable band shift by SDS-PAGE gel, even though Swi6 in these mutants is expected to retain phosphorylation at other sites. This was previously observed (Shimada et al. 2009) and is likely because the sequence context of a phosphorylated residue determines whether or not it will result in a bandshift (O’Donoghue and Smolenski 2022). When analyzed by flow cytometry, Δswi6 cells exhibit a silencing defect in which both the nucleation (green ON) and spreading (orange ON) reporters are expressed (Figure 1D). Conversely, wildtype swi6 cells show robust silencing of both reporters as we reported prior (Greenstein et al. 2018) (orange 11.1% ON, green 0.13% ON Figure 1E). Mutating only S46 and S52 to alanines (*swi6*^S46/52A^) largely phenocopies wildtype swi6 (orange 15% ON, green 0.06% ON Figure 1F). In contrast, mutation of serines at 18 and 24 (*swi6*^S18/24A^) resulted in the loss of spreading (orange 92.4% ON), while largely maintaining proper nucleation (green 2.65% ON) (Figure 1G, SFigure 1A-C). Restoring S18 and S24, while mutating the other 6 serines to alanines (*swi6*^S46/52/117-220A^) recovers much of the nucleation and spreading observed in wildtype, though with a modest silencing loss at orange (orange 26.4% ON, green 0.14% ON Figure 1H, SFigure 1D-F). Thus, S18 and S24 play a dominant role in regulating spreading, while other serines make a minor contribution. However, when only S46 and S52 are available (*swi6*^6S/A^), cells not only exhibit a loss of spreading (orange 94.3% ON) but also a modest loss of silencing at the nucleator (green 4.78% ON) (Figure 1I). This loss of silencing approaches but is not as severe as the deletion of ckb1, the gene encoding a crucial regulatory subunit of the CKII kinase: When we examined *Δckb1* in a separate experiment from the above, we found that the complete loss of Swi6 phosphorylation not only disrupts spreading (orange 94.4% ON versus 7.04% in the wildtype control, SFigure 1J-L), but also affects silencing at the nucleator (green 26.3%-38.0% ON versus 0.36% in the wildtype control). However, this defect is not nearly as severe as in *Δswi6* (Figure 1D), highlighting the role of Swi6 phosphorylation primarily in spreading. Overall, we interpret these results to indicate that NTE S18-52 phosphorylation contributes to regulating spreading, with S18/24 as major and 46/52 as minor contributors. Phosphorylation of serines in the Swi6 CD and hinge make a further minor contribution to Swi6’s overall silencing role, which is revealed only in the context of S18/24A. Given the greater loss of silencing revealed by Δckb1, we speculate that there are additional CKII target residues in Swi6, a notion confirmed by our *in vitro*Mass Spectrometry (see below), and that their phosphorylation contributes to Swi6’s silencing role at the nucleator.

### Serines 18 and 24 are required for the spreading of H3K9me3 but not H3K9me2

We next asked how phosphorylation of S18 and S24 contributes to the propagation of heterochromatic histone marks. We used chromatin immunoprecipitation followed by sequencing (ChIPseq) to address how levels of the heterochromatic marks, Histone 3 lysine 9 di- and trimethylation (H3K9me2/me3), are affected in the context of wildtype *swi6, swi6*^S18/24A^, and *Δswi6* in the MAT *ΔREIII* HSS background containing the “green” and “orange” reporters (Figure 2A). Consistent with prior work, we define H3K9me2 as the heterochromatin structural mark (Jih et al. 2017) and H3K9me3 as the heterochromatin spreading and silencing mark (Jih et al. 2017; Hall et al. 2002; Yamada et al. 2005). Following up on Figure 1, we initially examined the MAT locus (Figure 2B). Note that we cannot make definite statements about ChIP-seq signals over the “green” and “orange” reporters themselves, as the reporter cassettes harbor sequences that are duplicated 3-4 times in the genome (Greenstein et al. 2022), making ChIP-seq read assignment ambiguous. Overall, H3K9me2 levels at the MAT locus dropped significantly in *Δswi6*, consistent with prior work (Hall et al. 2002); however, *swi6*^S18/24A^ mutant maintained similar levels of H3K9me2 to wildtype swi6 (Figure 2B, top). Examining the distribution more closely, at the cenH nucleator, only *Δswi6* showed a minor decline of H3K9me2 in some regions. To the left of cenH, H3K9me2 levels decreased in *Δswi6* but not in *swi6*^S18/24A^ . To the right of cenH, H3K9me2 levels also severely declined in *Δswi6*, while in *swi6*^S18/24A^ they appear to drop moderately near mat3M, but recovered to wildtype levels at IR-R. When examining H3K9me3, we observed a different relationship: H3K9me3 patterns in *swi6*^S18/24A^ much more closely mirrored *Δswi6*. Specifically, to the left of *cenH*, H3K9me3 dropped to an intermediate level between wildtype and *Δswi6*, while on the right of cenH, H3K9me3 levels closely matched *Δswi6* (Figure 2B, bottom). Importantly, this behavior of H3K9me3 is consistent with our flow cytometry results (Figure 1), where silencing is largely unaffected at ‘green’ in *swi6*^S18/24A^, while ‘orange’ was expressed.

**Figure 2:**
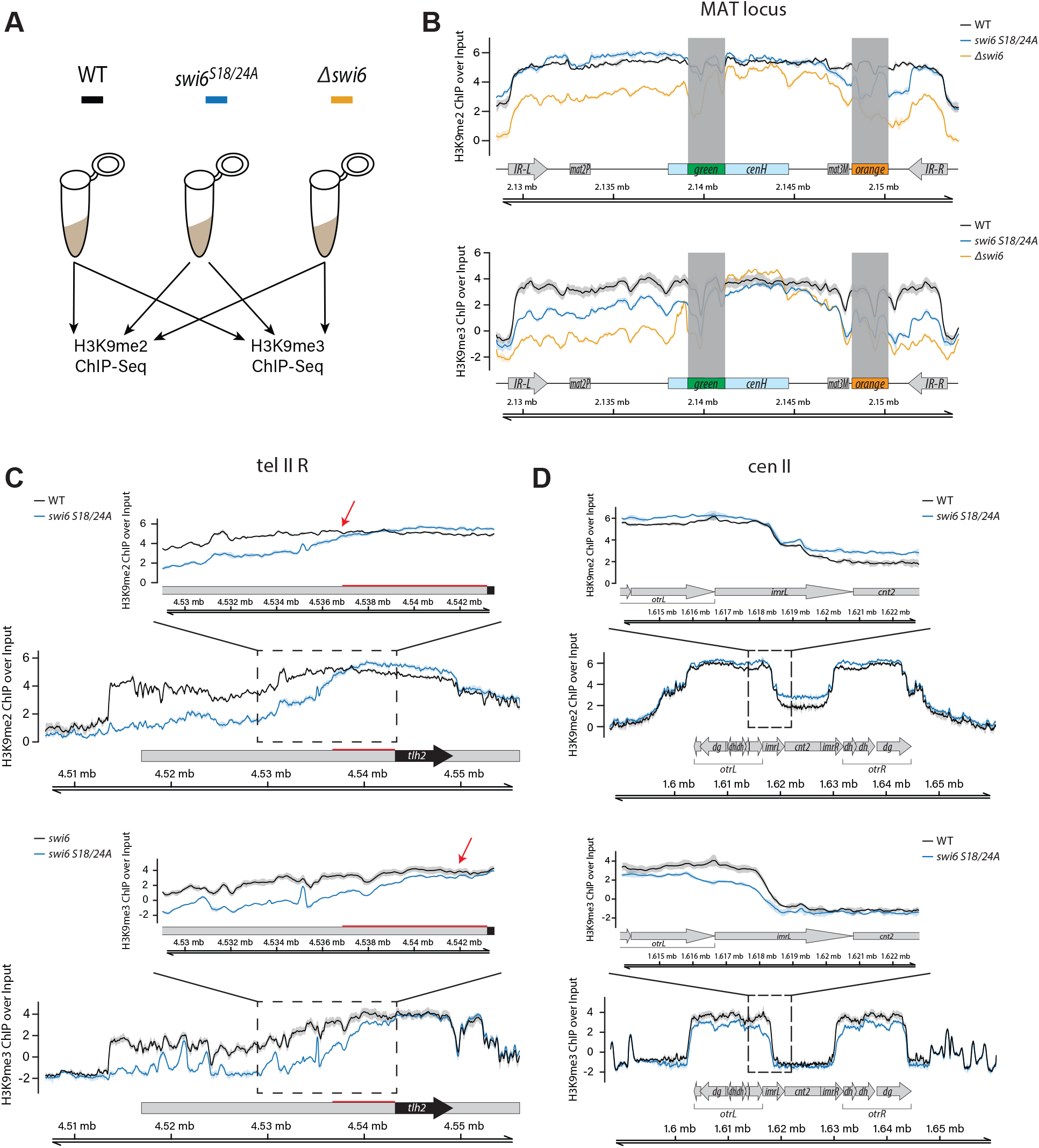
Conversion from H3K9me2 to H3K9me3 is compromised outside nucleation centers in S18 and S24 Swi6 mutants. **A**. Overview of the ChIP-seq experiments. **B-D**. ChIP-seq signal visualization plots. The solid ChIP/input line for each genotype represents the mean of three repeats, while the shading represents the 95% confidence interval. **B**. Plots of H3K9me2 (TOP) and H3K9me3 (BOTTOM) ChIP signal over input at the MAT *ΔREIII* HSS mating type locus for wildtype (black), *swi6*^S18/24A^ (blue), and *Δswi6* (gold). Signal over “green” and “orange” reporters are greyed out. Reads from these reporters map to multiple locations within the reference sequence, as all reporters contain control elements derived from the *ura4* and *ade6* genes. **C**. H3K9me2 (TOP) and H3K9me3 (BOTTOM) plots as in A. for subtelomere IIR for wildtype and *swi6*^S18/24A^. The red bar on the H3K9me2 plot indicates the distance from *tlh2* to where H3K9me2 levels drop in *swi6*^S18/24A^ relative to wildtype. Insets: a zoomed-in view proximal to *tlh2* is shown for H3K9me2 and me3. The red arrows in the insets indicate the point of separation of the 95% confidence intervals, which is significantly further telomere-proximal for H3K9me3. **D**. H3K9me2 (TOP) and H3K9me3 (BOTTOM) plots as in A. for centromere II for wildtype and *swi6*^S18/24A^. Insets: the left side of the pericentromere.

We wanted to further examine if the observation of H3K9me3 loss in *swi6*^S18/24A^ versus wildtype swi6 held for other genome regions. To examine this genome-wide, we binned the genome into 300bp bins and compared H3K9me2 and me3 accumulation between wildtype, *swi6*^S18/24A^, and *Δswi6*. We divided the K9me2/3 positive bins into regions we previously annotated as potential heterochromatin nucleation or spreading regions(Greenstein et al. 2022). We note that several of the features retrieved from PomBase as nucleators are likely internal spreading regions (Djupedal et al. 2009; Buscaino et al. 2013). Heat maps comparing the three genotypes (SFigure 2) show clearly the following trends: H3K9me3 is reduced in *swi6*^S18/24A^ versus wildtype in almost all spreading regions and moderately at nucleation-annotated sites, though only modestly at some bona fide nucleators such as *tlh1,2* and the 3′ of *cenH*. In contrast, H3K9me2 is elevated in almost all nucleators in *swi6S18/24A*, while in spreading regions, it is elevated in some but reduced in others. We note that, consistent with prior data, Δswi6 shows elevated H3K9me3 in several nucleators (Seman et al. 2023). We then analyzed some regions in more detail: We first analyzed the subtelomeric region (tel IIR). Consistent with the above analysis, we found that over the nucleation region *tlh2*, H3K9me2 levels are slightly elevated in *swi6*^S18/24A^, however, they then begin to drop ∼6.4 kb to the left of *tlh2* (Figure 2C, top, red bar, and arrow). Interestingly, H3K9me3 levels drop closer to the tlh2 nucleator than H3K9me2; the 95% confidence interval of wildtype and *swi6*^S18/24A^ separate at the left edge of tl*h2* (Figure 2C, bottom). This observation at the *tlh2* nucleator suggests the conversion of H3K9me2 to H3K9me3 is inhibited right as heterochromatin structures exit nucleation centers. We observed the same trend at the left subtelomere of chromosome I (tel IL, SFigure 3B). At the subtelomere, spreading distances outside nucleation sites are longer than at other loci; therefore, this loss of H3K9me3 just outside tlh2 manifests as an H3K9me2 spreading defect several kilobases downstream. This result is consistent with the requirement of Suv39/Clr4 methyltransferases to bind H3K9me3 for H3K9 methylation spreading (Al-Sady et al. 2013; Muller et al. 2016). We note that the left telomere of chromosome II contains no annotated nucleators in the published sequence. Hence, we could not observe the same trend there (tel IIL, SFigure 3C).

### Swi6 phosphorylation enables spreading

Second, we observed a similar defect in H3K9me3 spreading at the pericentromere (cenII), specifically, from the outer repeat (otr) into the inner repeat (imr) (Figure 2D, bottom versus top). However, the distances are likely too short from nucleation centers in otr to observe a resulting loss of H3K9me2 (Figure 2D). We note no distinguishable differences in H3K9me2 and H3K9me3 at *mei4*, a well-studied heterochromatin island (SFigure 3A).

Together, our ChIP-seq data show that *swi6*^S18/24A^ is deficient in the conversion of H3K9me2 to me3, especially outside nucleation centers. As evident for the subtelomere, this impairment results in the loss of silencing, and ultimately, the loss of H3K9me2 spreading which depends on H3K9me3 (Al-Sady et al. 2013; Jih et al. 2017).

### Swi6 phosphorylation increases oligomerization acting through, or in parallel to, known oligomerization surfaces

Next, we wanted to pinpoint the biochemical mechanisms that can account for the spreading defects in *swi6*^S18/24A^ (Figure 1G, Figure 2). HP1 oligomerization has been linked to spreading (Canzio et al. 2011). In turn, HP1’s intranuclear dynamics have been linked to how it engages chromatin (Cheutin et al. 2004; Hiragami-Hamada et al. 2011; Biswas et al. 2022; Williams et al. 2024). We thus probed if and how phosphorylation may impact these two properties of Swi6.

We used Size Exclusion Chromatography followed by Multi-Angle Light Scattering (SEC-MALS) to probe oligomerization. To produce phosphorylated Swi6 (pSwi6), we co-expressed Swi6 with Caesin Kinase II (CKII) in E. coli (Figure 3A). We used 2-dimensional Electron Transfer Dissociation Mass Spectrometry (2D ETD-MS) to identify which residues in pSwi6 are phosphorylated and used unphosphorylated Swi6 (unpSwi6) as a control (Figure 3B and SFigure 4A). We found that only pSwi6, and not unpSwi6, has detectable phosphorylated peptides. The residues phosphorylated in pSwi6 include several that were identified *in vivo* (Figure 1A, S18, S24, S46, S52, S117, S212, S220 but not S192) and some additional sites not previously identified (S43, S45, S165, S224, S227). This detection of additional CKII target sites is likely because of the higher sensitivity achieved in our 2D-ETD-MS experiments from purified protein: 1. 2D-ETD-MS better preserves phosphorylation sites compared to other methods and is highly sensitive. 2. Pure, in vitro-produced protein of high yield is likely to result in more detection events than *in vivo*-derived protein. Within the limits of our Mass Spectrometric analysis, we aimed to quantify the phosphorylation levels of *in vitro*purified pSwi6 at S18 and S24 in particular. We performed three independent analyses of separately injected pSwi6 material. The average of the three analyses indicates that the vast majority, ∼76%, of peptides are phosphorylated at S18 and/or S24 (SFigure 4A). We note that a larger fraction of S24 is persistently phosphorylated than S18. This may be an underestimate of the phosphorylated fraction, given the potential of losing phosphate groups in the peptide preparation.

**Figure 3:**
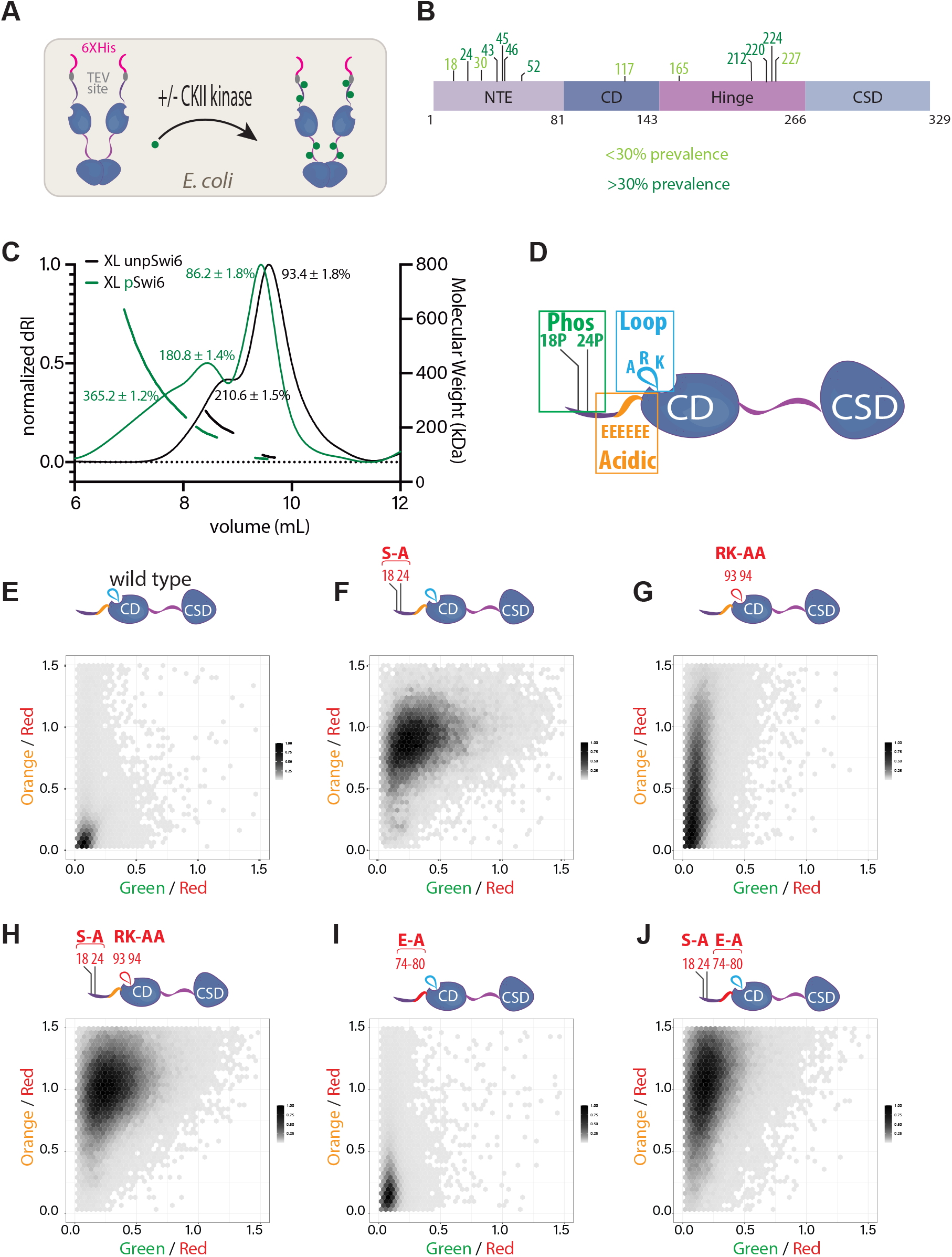
Swi6 phosphorylation increases oligomerization working through, or in parallel to, the ARK loop and NTE acidic stretch. **A**. Production of phosphorylated Swi6 (pSwi6) in *E. coli*. Casein Kinase II (CKII) is co-expressed with Swi6. After lysis and purification, the 6XHis tag is removed from the pSwi6 or unpSwi6 protein by TEV cleavage. **B**. Mass Spectrometry on pSwi6. Shown is a domain diagram of Swi6. Phosphorylation sites identified in pSwi6 by 2D-ETD-MS are indicated and grouped by detection prevalence in the sample. For prevalence of pS18 and/or p24 peptides, see SFigure 4A **C**. Size Exclusion Chromatography followed by Multi-Angle Light Scattering (SEC-MALS) on EDC/NHS cross-linked unpSwi6 (black) and pSwi6 (green). Relative refractive index signals (solid lines, left y-axis) and derived molar masses (lines over particular species, right y-axis) are shown as a function of the elution volume. The Swi6 concentration was 100µM. **D**. Overview of NTE and CD residues targeted for mutation in subsequent panels. S18/S24 phosphoserines, the CD ARK loop (R93AK94A; loopX), and the CD-preceding acidic stretch (E74-80A; acidicX). **E.-J**. 2-D Density hexbin plots examining silencing at nucleation ‘green’ and spreading ‘orange’ reporter, wildtype, S18/24A, loopX, and acidicX mutants and combinations thereof. Note this experiment was a fully separate run from Figure 1/SFigure 1.

SEC-MALS traces of uncrosslinked pSwi6 and unpSwi6 reveal both proteins are estimated to be of similar dimer mass, 90.8 kDa and 100.4 kDa respectively (SFigure 4B). However, pSwi6 elutes before Swi6, a trend similar to phosphorylated HP1α (Larson et al. 2017). There is also a small shoulder in the pSwi6 trace, indicating a minor fraction of higher-order oligomers (SFigure 4B, grey arrow). As previously published (Canzio et al. 2011), Swi6 crosslinking leads to the appearance of higher molecular weight species. We observed that crosslinked Swi6 and pSwi6 elute as apparent dimers (93.4 and 86.2 kDa, respectively) and tetramers (210.6 and 180.8 kDa, respectively) (Figure 3C). However, only pSwi6 additionally forms octamers (365.2kDa) and possibly even larger oligomers, as indicated by a broad shoulder (Figure 3C).

We wondered how S18/24 phosphorylation-mediated oligomerization relates to the known surfaces within Swi6 that promote higher-order oligomers (Canzio et al. 2011, 2013). Using our HSS system, we probed this relationship by constructing mutants combining S18/24A and the mutation of two previously characterized surfaces29: the ARK loop in the CD (R93A and K94A, termed “*swi6*^loopX^” ), and the acidic patch just preceding the CD (E74-80A, termed “*swi6*^acidicX^”, Figure 3D). The *swi6*^loopX^ mutant had a moderate spreading defect (Figure 3G, SFigure 5B). When combined with S18/24A, the *swi6*^loopX-S18/24A^ double mutant phenotype largely mimics that of *swi6*^S18/24A^ (Figure 3H, SFigure 5C) without significant further loss of ‘green’ (nucleation) or ‘orange’ (spreading) silencing. *swi6*^acidicX^ mutants only have a mild spreading phenotype (Figure 3I, SFigure 5D), and like swi6loopX-S18/24A, the double mutant *swi6*^acidicX-S18/24A^ experiences no significant loss of spreading or silencing beyond *swi6*^S18/24A^ alone (Figure 3J, SFigure 5E). We note that in this flow cytometry experiment, all strains, including wildtype, displayed slightly more fluorescence in ‘green’ than prior runs (compare Figure 3E, F to Figure 1E, G). Together, these results suggest that oligomerization of Swi6 is important for productive spreading. Further, they indicate that the known oligomerization contacts either work in parallel to, or require, phosphorylation at S18 and S24. However, because the *swi6*^S18/24A^ has a more severe spreading defect than the oligomerization surface mutants alone, S18 and S24 phosphorylation likely has additional molecular roles.

### Swi6 phosphorylation decreases nucleosome binding in vitro

We next quantified the binding of pSwi6 to its target substrate (Figure 4A). We first measured the association of pSwi6 with H3K9me0 and H3K9me3 peptides by fluorescence polarization (Figure 4B). pSwi6 binds to H3K9me0 and H3K9me3 peptides with affinities (Kd) of 227.4µM and 2.45µM, respectively, revealing a ∼93X specificity for H3K9me3 (Figure 4D). While we could not determine the H3K9me0 peptide Kd for unpSwi6, the Kd for the H3K9me3 peptide was 8.17 µM (Figure 4B, D). Previously, the specificity for unpSwi6 was reported at ∼130X (Canzio et al. 2011), thus indicating little difference in H3K9me3 peptide specificity between the two proteins. We note that consistent with previous reports on total cellular Swi6 (Jih et al. 2017), recombinant pSwi6 also shows a ∼2.2X preference for H3K9me3 versus H3K9me2 peptides (SFigure 4C).

**Figure 4:**
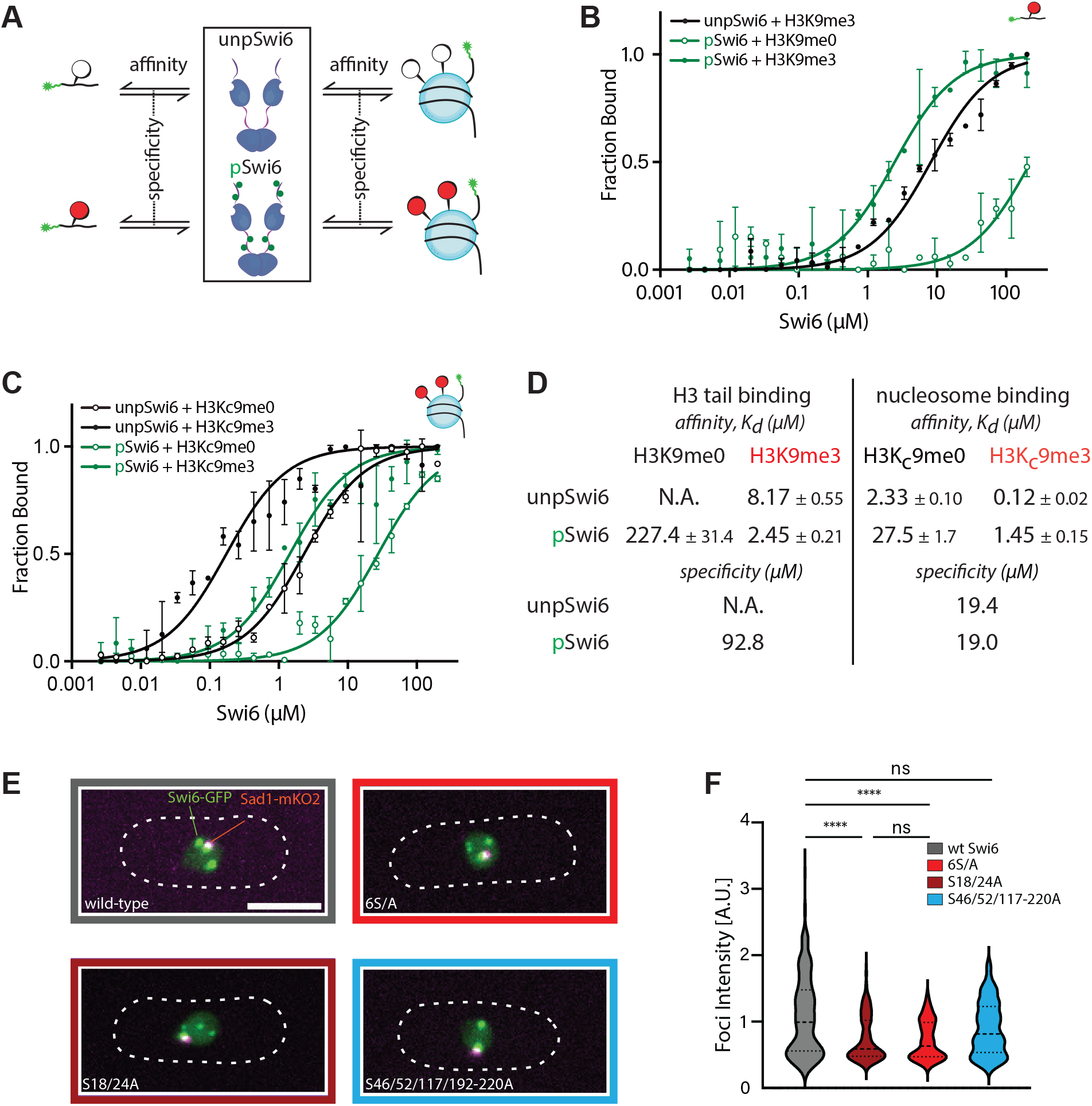
Swi6 phosphorylation decreases nucleosome affinity without affecting specificity. **A**. Overview of fluorescence polarization (FP) experiments with fluorescein (star)-labeled H3 tail peptides (1-20) and nucleosomes to assess pSwi6 and unpSwi6 substrate affinity and specificity. **B**. FP of H3K9me0 (open circles) and H3K9me3 (filled circles) tail peptides with pSwi6 (green) or unpSwi6 (black). The binding affinity was too low to be fit for unpSwi6 and H3K9me0 peptides. **C**. FP with H3K9me0 (open circles) or H3Kc9me3 (MLA, filled circles) mononucleosomes. Fluorescein (green star) is attached by a flexible linker at one end of the 147 bp DNA template. For B.&C., the average of three independent fluorescent polarization experiments for each substrate is shown. Error bars represent standard deviation. **D**. Summary table of affinities and specificities for B. and C. **E**. Representative maximum projection live microscopy images of indicated Swi6-GFP/ Sad1-mKO2 strains. **F**. Analysis of signal intensity in Swi6-GFP foci in indicated strains. Wt Swi6, n=242; Swi6^S18/24A^, n=251; Swi6^S18/24/117-220A^ (6S/A), n=145; Swi6^S46/52/117-220A^, n=192. n, number of foci analyzed.

We next probed how phosphorylation affects nucleosome binding. We performed fluorescence polarization with fluorescently labeled nucleosomes that are either unmethylated (H3K9me0) or trimethylated (H3Kc9me3) (Simon et al. 2007),(Canzio et al. 2011). Phosphorylation had no impact on the specificity for the H3K9me3 mark, consistent with the peptide observation (19.4X, vs. 19X for unpSwi6 or pSw6, respectively, Figure 4C, D).

However, we observe a 12X difference in affinity to the nucleosome overall between pSwi6 and Swi6 (Figure 4D). The H3Kc9me3 nucleosome affinity is 0.12 µM and 1.45 µM for unpSwi6 and pSwi6, respectively, while the H3K9me0 affinity is 2.33 and 27.5 µM, respectively. We note the affinity of pSwi6 to the H3Kc9me3 nucleosome is similar to its affinity to the H3K9me3 peptide, binding only 1.7X tighter to the H3Kc9me3 nucleosome (1.45µM vs. 2.45µM). Instead, and consistent with previous results, unpSwi6 binds 68X more tightly to the nucleosome than to the tail (8.17 µM for the H3K9me3 tail versus 0.12µM for H3Kc9me3), which is thought to arise from additional contacts beyond the H3 tail on the nucleosome.

Why would a 12X lower affinity towards the nucleosome substrate be advantageous for pSwi6’s function in spreading (Figure 1,2)? In the literature, the cellular abundance of Swi6 is measured at 9000-19,400 molecules per cell (Carpy et al. 2014; Sadaie et al. 2008). The estimated fission yeast nuclear volume of ∼7mm3 (Lemière et al. 2022; Neumann and Nurse 2007) then yields an approximate intranuclear Swi6 concentration of ∼2.1 -4.6µM. Given our measured nucleosome Kds (Figure 4D), the intranuclear concentration of unpSwi6 would theoretically be above its Kd for both H3K9me0 and me3 nucleosomes. The concentration of pSwi6 would exceed its Kd for H3Kc9me3 but be significantly below (∼10X) its Kd for H3K9me0 nucleosomes. We cannot assume the same fraction of bound nucleosome from *in vitro*measurements applies *in vivo*, because nucleosome concentrations in the cell (∼10µM based on accessible genome size and average nucleosome density (Lantermann et al. 2010; Godde and Widom 1992)) greatly exceed what is used in a binding isotherm. We can use a quadratic equation (Jarmoskaite et al. 2020) (see methods) appropriate for these *in vivo* regimes instead of a typical Kd fit to estimate the fraction bound. As only 2% of the *S. pombe* genome is heterochromatic, we approximate the total nucleosome concentration (10µM) to reflect unmethylated nucleosomes. The small, methylated nucleosome pool will mostly be bound by Swi6 irrespective of the phosphorylation state. However, we estimate that only 5% of unmethylated nucleosomes would be bound by pSwi6, while this would be ∼16% for unpSwi6. At the high end of the Swi6 concentration estimate, this fraction bound would increase to 30% of unmethylated nucleosomes. Further, we expect enhanced oligomerization of pSwi6 on heterochromatin to reduce the free Swi6 pool (see discussion). Therefore, we predict that the main function of phosphorylation is to limit the partitioning of Swi6 into the unmethylated pool, confining it to heterochromatin.

### Phosphorylation directs Swi6 into heterochromatin nucleators away from euchromatic sites

One test of the above prediction would be altered localization of wildtype and phosphorylation defective Swi6 versions in the fission yeast nucleus. Across species, HP1 homologs have been shown to localize into heterochromatic foci *in vivo* and form LLPS droplets *in vitro*(Larson et al. 2017; Strom et al. 2017; Sanulli and J Narlikar 2020; Cheutin et al. 2004). Specifically, phosphorylation of the NTE in human HP1α is one driver of heterochromatin foci formation (Larson et al. 2017; Hiragami-Hamada et al. 2011). We investigated whether the loss of phosphorylation sites that impair heterochromatin spreading (Figure 1, 2) impacted partitioning between heterochromatin foci and regions outside these foci, likely representing H3K9 unmethylated nucleosomes. We C-terminally tagged wildtype swi6 and phospho-serine mutants at the native locus with super fold-GFP (Swi6-GFP), as an N-terminal tag disrupts Swi6 dimerization and oligomerization (Canzio et al. 2013). We crossed these strains into a background containing sad1:mKO2, a spindle pole body (SPB) marker (SFigure 6A). We chose this background as Sad1 denotes the position of pericentromeric heterochromatin (Hou et al. 2012; Barrales et al. 2016) and can help orient other heterochromatin sites relative to it. We examined the following SF-GFP tagged mutant variants: *swi6*^S18/24A^, *swi6*^S46/52A^, *swi6*^S46/52/117-220A^ (S18/S24 available), and *swi6*^6S/A^ (Figure 1G-I) and imaged these strains by confocal microscopy (Figure 4E, SFigure 6B). Largely, these mutations do not impact either Swi6 accumulation (SFigure 6C), nuclear foci number (SFigure 6D), or position of the foci relative to the SPB (Al-Sady et al. 2016) (SFigure 6E, F).

We next quantified the accumulation of Swi6-GFP in foci (Figure 4E, F). Unlike foci number or spatial arrangement, the average foci intensity for Swi6-GFP strains carrying the S18/24A mutations is significantly decreased relative to wildtype Swi6-GFP (Figure 4F), while the nucleoplasmic signal increases. Because total Swi6GFP levels do not change in these mutants (SFigure 6C), this result indicates that Swi6^S18/24A^-GFP and Swi6^6S/A^ -GFP molecules partition away from heterochromatin foci. This finding is consistent with our prediction based on our *in vitro*measurements and implies that Swi6 molecules that cannot normally be phosphorylated partition onto unmethylated nucleosomes.

To more specifically determine if, and to which genomic locations, phospho-mutant Swi6 molecules are redistributed, we performed two ChIP-seq experiments: 1. With FLAG-tagged versions of wildtype *swi6, swi6*^S18/24A^ *swi6*^S18/24/117-220A^ (*swi6*^6S/A^), using an anti-FLAG antibody. This allowed us to track if and how the phospho-mutant distribution changes versus wildtype. 2. With untagged wildtype and *swi6*^S18/24A^ strains, using a phospho-serine antibody specific for phosphorylation at S18 and S24 (anti-pS18-pS24, see Figure 6A below). This allowed us to determine where pS18 and/or pS24-phosphorylated Swi6 accumulates. For accurate quantitation of signal across strains, we used a spike-in control (see methods, spike-in adjusted signal quantified as counts per million, cpm). To broadly examine Swi6 distribution, we compared accumulation of wildtype FLAG-Swi6, FLAG-Swi6^S18/24A^, and FLAG-Swi6^6S/A^ in a heatmap, examining all genomic H3K9me-positive loci, divided into nucleation sites and spreading zones, as above (SFigure 2)(Greenstein et al. 2022). We noticed a decline in accumulation of FLAG-Swi6^S18/24A^ and FLAG-Swi6^6S/A^ versus wildtype at many nucleation sites (Figure 5B), consistent with the loss in heterochromatin foci (Figure 4E, F). We observed an increase for FLAG-Swi6^S18/24A^ at some sites, notably in the pericentromere, although some of those are likely internal spreading zones (see below). When we compared “spreading sites” we observed declines at some locations (MAT locus, some subtelomeres) but also increases (tel IIR, inner most repeats, and the central cores, Figure 5C). These results suggest situations where FLAG-Swi6^S18/24A^ and/or FLAG-Swi6^6S/A^ may be binding to nucleosomes either in the process of, or prior to, methylation.

**Figure 5:**
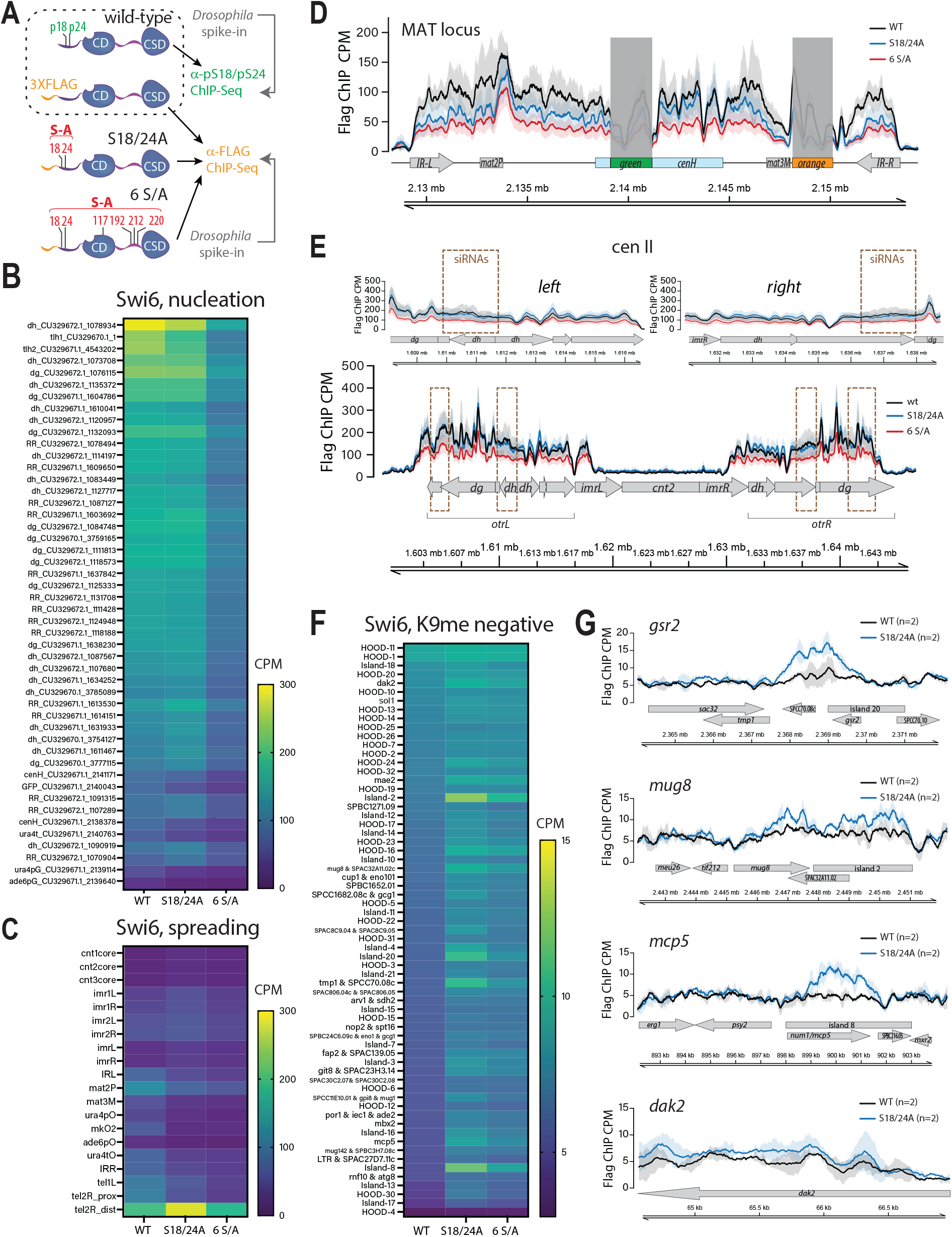
Swi6 phosphorylation focuses Swi6 onto heterochromatin nucleation sites and away from euchromatin. **A**. Schematic of Swi6 ChIP-seq experiments. The anti-pS18-pS24 experiment was carried out with wildtype *swi6* (and *swi6*^*S18/24A*^ as a negative control, see **SFigure 8**). For the anti-FLAG experiments, the endogenous *swi6* locus was 3XFLAG tagged in the context of wildtype *swi6* (black), *swi6*^*S18/24A*^ (blue), or *swi6*^*S18/24/117-220A*^ *(6S/A*, red). For quantitative normalization, ChIP reactions were supplemented with *Drosophila* chromatin spike-in. **B**. Heatmaps of spike-in normalized FLAG ChIP-seq signal (in Counts Per Million, CPM) for *swi6, swi6*^*S18/24A*^, or *swi6*^*6S/A*^ at features previously classified as nucleators(Greenstein et al. 2022). **C**. As in B. but for features classified as regions of H3K9me2 spreading(Greenstein et al. 2022). **D**. FLAG ChIP-seq signal visualization plot for the MAT *ΔREIII* HSS mating type locus. The solid ChIP/input line for each genotype represents the mean of three repeats, while the shading represents the 95% confidence interval. **E**. As in D. but for cen II with zoom-in of the left and right pericentromeric region. The brown dashed boxes indicate siRNAi-generating centers as mapped in [(Djupedal et al. 2009)]. **F**. Heatmap as in B. and C., but for H3K9me2 negative regions. **G**. ChIP-seq signal visualization plots for H3K9me2 negative regions. The solid ChIP/input line for each genotype represents the mean of two repeats, while the shading represents the 95% confidence interval. A high signal replicate was removed for both genotypes. ChIP-seq signal visualization for all three genotypes with all three repeats in SFigure 7C.

**Figure 6:**
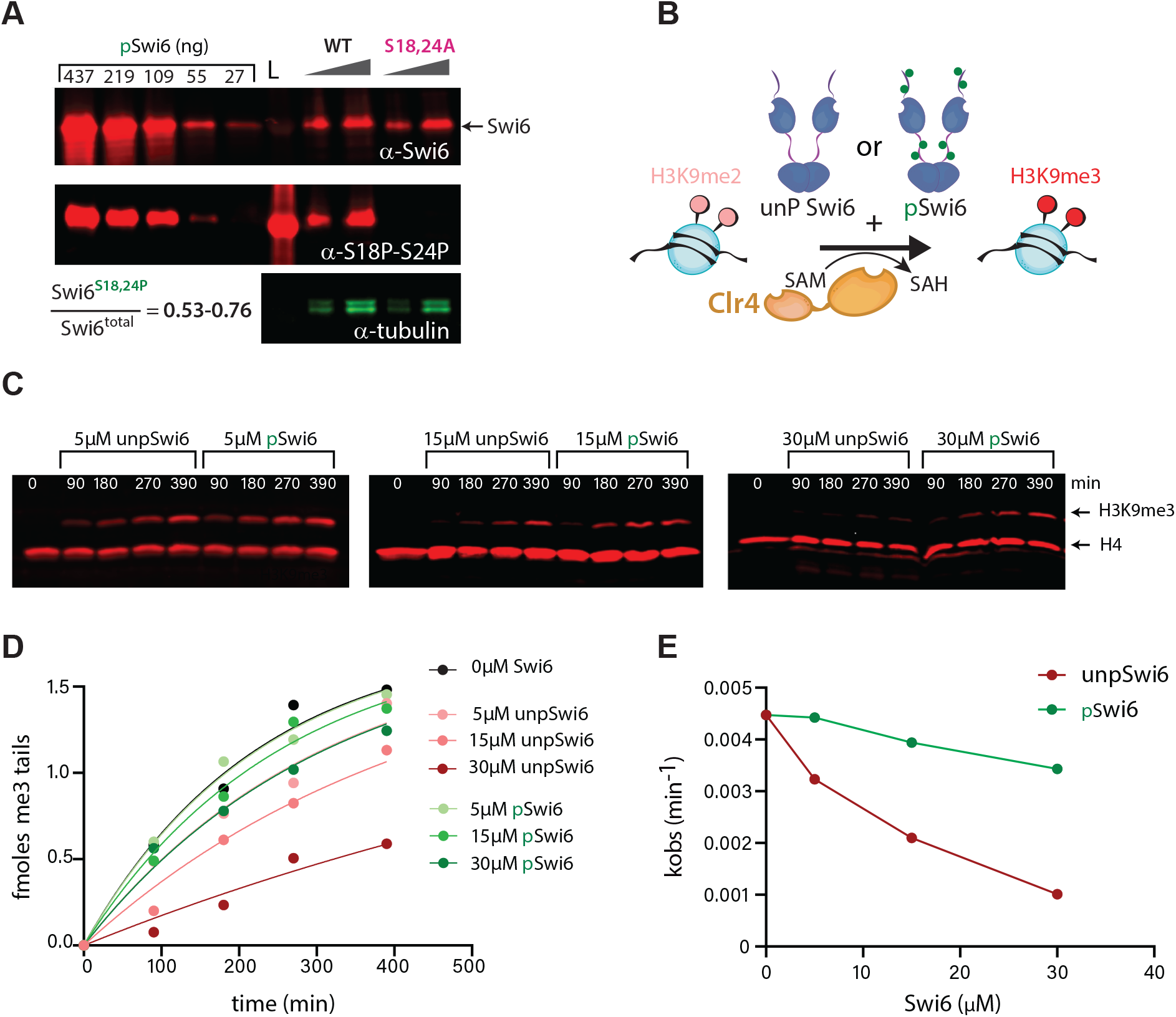
Swi6 phosphorylation mitigates inhibition of the Clr4-mediated conversion of H3K9me2 to H3K9me3. **A**. Most Swi6 molecules in the cell are phosphorylated at S18 and S24. Quantitative western blots against total Swi6 and phosphorylated Swi6 at S18 and/or S24. A standard curve of pSwi6 isolated as in Figure 3 is included in both blots. Total protein lysates from wildtype *swi6* and *swi6*^S18/24A^ strains were probed with a polyclonal anti-Swi6 antibody (α-Swi6) or an antibody raised against a phosphorylated S18/S24 peptide (α-pS18-pS24). α-tubulin was used as a loading control. One of two independent experiments is shown. L; ladder. Total fraction of Swi6 phosphorylated *in vivo* at S18 and/or S24 is adjusted by the prevalence of phosphorylation in the *in vitro* produced standard (∼0.76, see SFigure 4A) **B**. Experimental scheme to probe the impact of Swi6 on H3K9 trimethylation. **C**. Quantitative western blots on the time-dependent formation of H3K9me3 from H3K9me2 mononucleosomes in the presence of pSwi6 or unpSwi6 under single turnover conditions. The same blots were probed with α-H3K9me3 and α-H4 antibodies as a loading and normalization control. **D**. Single exponential fits of production of H3K9me3 tails over time for indicated concentrations of unpSwi6 or pSwi6. **E**. plot of the observed single turnover rate constant measured from exponential fits (kobs) against the Swi6 concentration in µM.

When we examine cen I (SFigure 7A) and cen II (Figure 5E) more closely, we noticed FLAG-Swi6S18/24A accumulates less than wildtype within previously mapped siRNA generating centers (Djupedal et al. 2009). However, just outside these centers, Swi6S18/24A levels rise to levels above wildtype (Figure 5E), coinciding with reduced H3K9me3 (Figure 2D). These locations likely represent internal spreading zones within the pericentromere. Swi66S/A is broadly depleted at cen I & II.

In contrast, the MAT locus (Figure 5D), tel IIL (SFigure 7B, bottom ), and tel IIR (SFigure 7B, top) show broadly reduced accumulation of FLAG-Swi6^S18/24A^ and Swi6^6S/A^ in dense H3K9me2/3 areas (Figure 2B, SFigure 3C, Figure 2C, respectively). Tel IIL shows slight, but consistent, FLAG-Swi6^S18/24A^ accumulation above wildtype in the adjacent area with low H3K9me2/3 (SFigure 7B, bottom), indicating Swi6 redistribution to tel IIL-adjacent euchromatin. Conversely, when we examined regions that are filtered to be H3K9me2-negative in our data and are located in euchromatin, we find several sites with significant accumulation of FLAG-Swi6^S18/24A^ and/or FLAG-Swi6^6S/A^ above wildtype (Figure 5F). Interestingly, this is most apparent in regions that can be facultatively methylated, such as HOODs (Yamanaka et al. 2013), islands (Zofall et al. 2012), or regions that can become heterochromatic when antagonists mst2 and epe1 are deleted (Wang et al. 2015). Because the FLAG-Swi6 ChIP-Seq is much noisier than H3K9me2/3, especially in the lower signal regime of euchromatin, we wanted to ensure that the euchromatic region differences between genotypes are not driven by repeats with higher signal. We thus plotted example regions either excluding one high-signal replicate of FLAG-Swi6 and FLAG-Swi6^S18/24A^ (Figure 5G) or including all replicates from all genotypes (SFigure 7C). In either case, we see clearly that FLAG-Swi6^S18/24A^ accumulates above wildtype at several H3K9me2 negative loci, thus binding unmethylated nucleosomes.

Our experiment with the anti-pS18-pS24 antibody indicated first, that the antibody is highly specific (see SFigure 8B). Second, when compared to wildtype FLAG-Swi6, there are no significant differences in accumulation between the overall Swi6 pool and the pS18-pS24 pool (SFigure 8F), indicating that the majority of the total Swi6 protein is likely phosphorylated at these residues (see below). Third, although pS18 and/or pS24 Swi6 generally overlaps with wildtype FLAG-Swi6, we observed that pS18 and/or pS24 Swi6 is overrepresented relative to wildtype FLAG-Swi6 at cen II siRNA generating centers (SFigure 8E). Additionally, there is a gradual depletion of relative pS18-pS24 signal compared to wildtype FLAG-Swi6 in the spreading zone of tel IIR (SFigure 8D). We note that comparisons of ChIP-seq conducted with different antibodies (see methods) have limitations, which we do not face with the above wildtype FLAG-Swi6 versus mutant Swi6 (Figure 5 and SFigure 7). Overall, these data are consistent with the notion that S18 and/or S24 phosphorylation focuses Swi6 at nucleators.

### Swi6 phosphorylation facilitates the conversion of H3K9me2 to me3 by Clr4

As unmethylated nucleosomes are Clr4’s substrates, another prediction emerges. Since unpSwi6 is more likely to bind unmethylated nucleosomes, Swi6 phosphorylation mutants may interfere with Clr4 substrates, which could explain the defect in H3K9me2 to me3 conversion in *swi6*^S18/24A^ (Figure 2), the slowest transition catalyzed by Clr4 (Al-Sady et al. 2013). For Swi6 phosphorylation to prevent the conversion of H3K9me to me3, the Swi6 cellular pool would have to be mostly in the phosphorylated state. To test this, we asked what fraction of Swi6 molecules in the cell are phosphorylated at S18 and S24. We addressed this question by a quantitative western blot approach, using two antibodies: a polyclonal Swi6 antibody (Canzio et al. 2013) to detect all Swi6 molecules and our pS18-pS24 specific antibody (top blot vs. bottom blot, respectively, Figure 6A, also see SFigure 8A-C). A standard curve of recombinant pSwi6 allowed us to quantify the total pool of Swi6 molecules vs. those phosphorylated at S18 and S24. The swi6S18/24A mutant control shows these phospho-serine antibodies are also specific under Western blot conditions (Figure 6A). Our data indicate after adjustment to the degree of phosphorylation in our *in vitro*standard (∼76%, SFigure 4A) that majority of cellular Swi6, ranging from 53% - 76% in biological replicate 1 and 2, respectively, is phosphorylated at S18 and/or S24 (Figure 6A and SFigure 9A). The finding that the majority of the Swi6 pool exists as pS18 and/or pS24 is corroborated by the overall overlap in the FLAG versus anti-pS18-pS24 ChIP-Seq (Figure 8F).

We next tested if Swi6 phosphorylation directly impacted the ability of Clr4 to produce H3K9me3. We incubated pSwi6 or unpSwi6 with Clr4 and monitored the conversion of the H3K9me2 substrate to H3K9me3 under single turnover conditions(Al-Sady et al. 2013) (Figure 6B). pSwi6 shows an *in vitro*preference for H3K9me3 versus H3K9me2 peptides (SFigure 4C), suggesting that phosphorylation may partition Swi6 towards H3K9me3 versus me0, but also, to some extent, towards H3K9me3 versus me2.

We observed that the presence of unpSwi6 inhibits the conversion of H3K9me2 to H3K9me3 in a concentration-dependent manner, but that this inhibition is significantly alleviated by pSwi6 (Figure 6C and SFigure 9B,C). Note we observe inhibition at the lowest concentration, 5µM, which is near the estimated *in vivo* concentration of Swi6. When normalizing to H4 and fitting H3K9me3 production to observed exponential single turnover rates (kobs), Clr4 methylation rates are significantly slowed in the presence of unpSwi6, while pSwi6 reduces this inhibition (Figure 6D,E).

While these data could explain the H3K9 trimethylation defect we observed for *swi6*^S18/24A^ (Figure 2), our in vitro-produced pSwi6 is phosphorylated at multiple residues. Given that pS18 and pS24 only represent around 1/6 of the detected phosphorylation sites (SFigure 4A), we cannot necessarily conclude whether the biochemical phenotypes we observe depend on S18 and S24 phosphorylation. To examine this, we expressed and purified a phospho-mutant protein, pSwi6^S18/24A^, in which S18 and S24 are mutated to alanines and co-expressed it with CKII (Figure 7A). pSwi6^S18/24A^ is still phosphorylated to a similar degree as pSwi6, which is apparent by the similar gel migration shift observed for both proteins (SFigure 10A, B). Upon phosphatase treatment, pSwi6^S18/24A^ and pSwi6 adopt the same migration pattern as unpSwi6 (SFigure 10A, B). 2D ETD-MS analysis of pSwi6^S18/24A^ additionally confirmed a similar phosphopeptide pattern to pSwi6, though with small changes in phosphopeptide prevalence (SFigure 10C).

**Figure 7:**
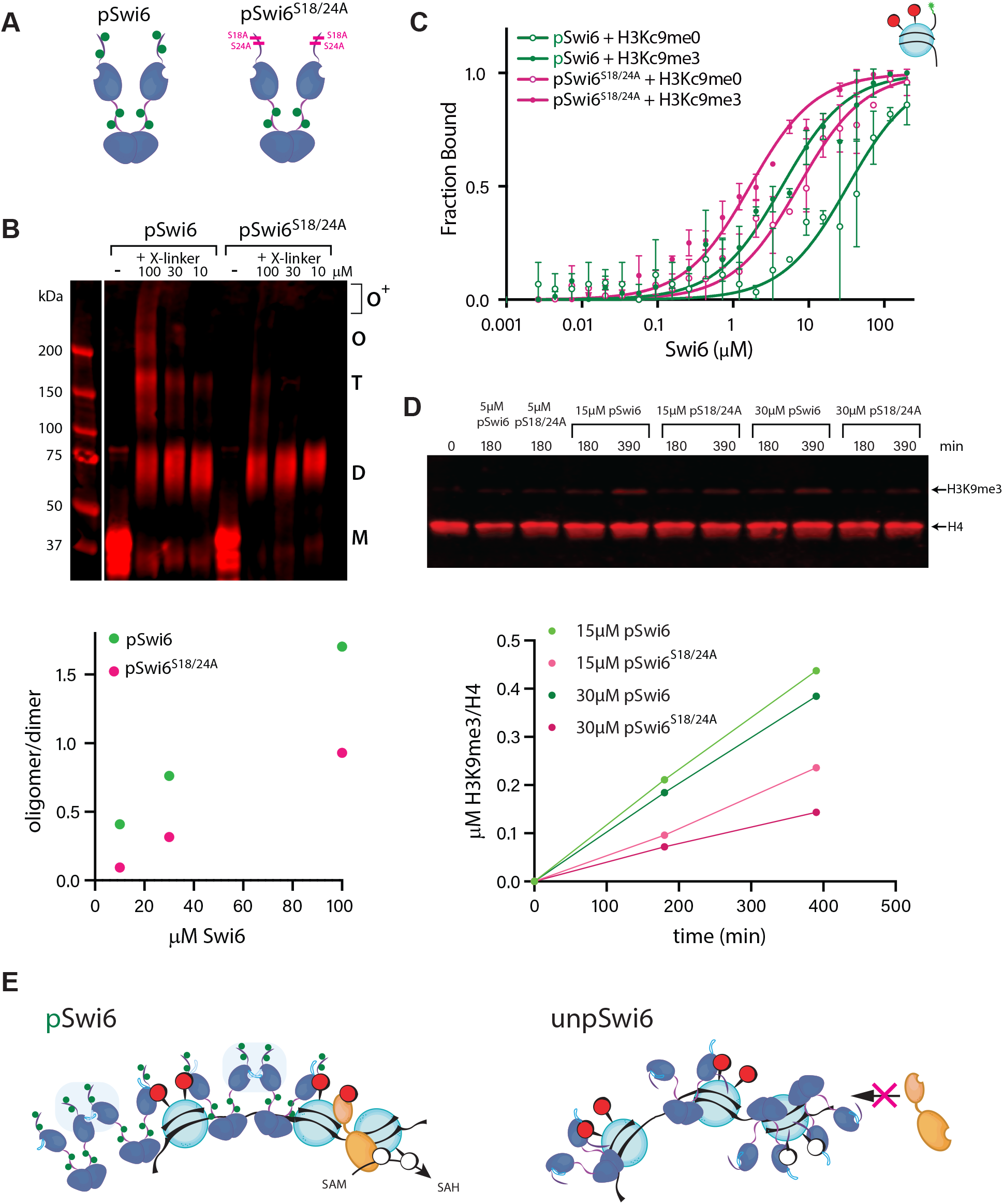
S18 and/or S24 phosphorylation contributes to pSwi6’s biochemical behaviors. **A**. Schematic of phosphorylated Swi6 molecules used in this Figure. **B**. pSwi6^S18/24A^ is defective in oligomerization. pSwi6 or pSwi6^S18/24A^ was crosslinked or not (-) at indicated concentrations, separated on SDS-PAGE, and probed with a polyclonal anti-Swi6 antibody. M, monomer; D, dimer; T, tetramer; O, octamer; O^+^, higher molecular weight species. Below: Quantification of oligomer signal divided by dimer signal for crosslinked species. **C**. FP with H3K9me0 (open circles) or H3Kc9me3 (MLA, filled circles) mononucleosomes as in Figure 4C, with pSwi6 (green) and pSwi6^S18/24A^ (magenta). Error bars represent the standard deviation of three repeats. Relative dissociation constant (Kd) values in SFigure 10D. **D**. Quantitative western blots on the time-dependent formation of H3K9me3 from H3K9me2 mononucleosomes in the presence of pSwi6 or pSwi6^S18/24A^, as in Figure 6C. Quantification of the signal below. Note that the reactions were not fast enough in this experiment to derive a single exponential observed rate. **E**. Model of the impact of pSwi6 on Clr4 activity. Left: pSwi6 does not engage with H3K9me0 nucleosomes, clearing the substrate for Clr4, and has reduced interactions with the nucleosome core. The pS18/pS24 NTE releases the ARK loop (light blue), allowing CD-CD contacts. Right: unpSwi6 binds H3K9me3 and me0 nucleosomes, occluding Clr4 access. The S18/S24 unphosphorylated NTE blocks the ARK from engaging CDs and additionally may contribute to increased nucleosome affinity by contacting DNA or octamer.

This pSwi6^S18/24A^ mutant has a strongly decreased propensity to oligomerize, compared to pSwi6 (Figure 7B) and has increased affinity towards both the H3K9me0 and Kc9me3 nucleosomes, 4.5 and 2.6X, respectively (Figure 7C, SFigure 10D). This result is consistent with pS18 and/or pS24 acting to modulate Swi6’s chromatin affinity. However, since the change in affinity for pSwi6S18/24A is less than the 12X loss observed for unpSwi6 vs. pSwi6, this implies that other phosphoserines also contribute to lowering nucleosome affinity. This results also accounts for the somewhat more severe *swi6*^6S/A^ phenotype (Figure 1G vs I).

We next checked whether pSwi6^S18/24A^ also inhibits the H3K9me2 to H3K9me3 conversion. Indeed, we see that just the loss of phosphorylation at S18 and S24 leads to a decrease in trimethylation (Figure 7D), likely due to the increased nucleosome affinity we observe above (Figure 7C, SFigure 10D). We note that in this set of experiments, trimethylation proceeded overall at a slower rate.

This data suggests a model whereby Swi6 NTE phosphorylation, particularly at S18 and/or S24, partitions Swi6 away from binding the unmethylated substrate of Clr4 *in vivo*, which is likely enhanced by increased Swi6 oligomerization at heterochromatin sites. Together, both reduced affinity and oligomerization mechanisms promote the H3K9me3 spreading reaction.

## Discussion

Previous work (Shimada et al. 2009) identified key Swi6 phosphoserines that regulate transcriptional gene silencing. In this work, we find that Swi6 phosphoserines 18 and 24 are required for heterochromatin spreading, but not nucleation (Figure 1). Swi6 phosphorylation promotes oligomerization (Figure 3) and tunes Swi6’s overall chromatin affinity to a regime that allows Clr4 to access its substrate (Figure 4), facilitating the conversion of dimethyl H3K9 to the repressive and spreading-promoting trimethyl H3K9 state (Figure 6). This modulation of chromatin affinity *in vivo* restricts Swi6 to heterochromatin foci (Figures 3, 5, SFigures 6, 7), which suggests that phosphorylation of HP1 molecules may be required for their concentration into the heterochromatic compartment. Three central themes emerge from this work:

### Swi6 phosphorylation decreases chromatin affinity, but not specificity

Phosphorylation is known to regulate HP1’s affinity with itself (Larson et al. 2017), DNA (Hiragami-Hamada et al. 2011; Nishibuchi et al. 2014), and chromati n(Nishibuchi et al. 2014; Hiragami-Hamada et al. 2011), but in manners that are homolog-specific. For example, phosphorylation in the NTE of HP1α induces LLPS, but not for HP1a in Drosophila, where phosphorylation instead regulates chromatin binding (Strom et al. 2017; Zhao et al. 2001; Zhao and Eissenberg 1999). Underlying this may be that CKII target sequences are not conserved across HP1s, for example, HP1α is phosphorylated in a cluster of 4 serines at the NTE (S11-14) (LeRoy et al. 2009), HP1a only at S15 in the NTE, and S202 C-terminal to the CSD (Zhao et al. 2001), whereas we report here Swi6 is phosphorylated by CKII in the NTE, CD, and hinge (Figure 3B).

The impact on nucleosome specificity is different across species. Our data here shows that phosphorylation of Swi6 does not affect its specificity for both H3K9me0 and H3Kc9me3 nucleosomes (Figure 4C, D), but phosphorylation of HP1α and HP1a was reported to increase its specificity for H3K9me3 nucleosomes (Larson et al. 2017; Nishibuchi et al. 2014). Instead, Swi6 phosphorylation decreases overall nucleosome affinity for unmethylated and H3Kc9me3 nucleosomes to a similar degree, 11.8X and 12X respectively, in contrast to HP1α (Nishibuchi et al. 2014, 2019). What may explain these differences? Internal interactions between the NTE, CD, hinge, and CSD work together to drive nucleosome binding (Canzio et al. 2013; Sanulli et al. 2019). We speculate that these domain interactions are differentially impacted by 1. the unique phosphorylation patterns in different HP1 orthologs (see above) and 2. divergence in Swi6 amino acid sequence and size of the NTE (Bensaha et al. 2025) and hinge that harbor most CKII target sites. Both these differences result in unique outcomes with respect to nucleosome specificity and affinity in different HP1 orthologs. Why might nucleosome affinity drop in the case of Swi6? One possibility is overall charge repulsion: While the pI of Swi6 is already negative (5.6), it is possible that the phosphorylation pattern on Swi6 locally increases net negative charge in a way that repels DNA and reduces affinity, consistent with prior reports on HP1 nucleic acid binding (Nishibuchi et al. 2014). This view is also supported by the observation that additional reduction in charge by serine to alanine mutations or lack of phosphorylation (*swi6*^6S/A^ versus *swi6*^S18/24A^ *in vivo*, pSwi6^S18/24A^ versus unpSwi6 *in vitro*Figures 4, 5, and 7) can have more severe effects than loss of phosphorylation at S18/S24 alone. Alternatively, rather than a net negative charge increase directing affinity loss, it may be the case that the NTE around S18/S24 makes specific contacts with the nucleosome. Cross-linking mass spectrometry studies indicate that the NTE of HP1α (Sokolova et al. 2024) and Swi6 (Sanulli et al. 2019) make contacts with multiple regions of histones including H2B, the C-terminus of H2A.Z (HP1α)/H2A (Swi6), and the core (HP1α) and tail (Swi6) of H3, among other contacts. NTE phosphorylation may specifically decrease these contacts, leading to detachment from the nucleosome core. The loss of this contact then decreases nucleosome affinity and may promote a change in Swi6 activity, such as oligomerization (Figure 3). This is further supported by the disproportionally stronger phenotype of mutants that carry S18/24A mutations versus any mutant with equal number of phosphosite-mutations that leave S18/S24 intact (Figure 1, Figure 4D,F, see discussion below).

This overall decrease in affinity partitions pSwi6 in a different way than unpSwi6, restricting access of pSwi6 to chromatin inside nuclear foci and away from euchromatic loci. This is supported by our imaging (Figure 4, SFigure 6) as well as our FLAG-Swi6 and pS18-pS24 ChIP-Seq data (Figure 5, SFigure 8). The redistribution to euchromatin is not even and some specific sites seem to accumulate phospho-mutant Swi6 more than others (Figure 5G, SFigure 7C). Since ∼105 euchromatic nucleosomes compete for approximately ∼1-2×104 Swi6 molecules not bound at heterochromatin, we speculate that euchromatic regions most able to compete for Swi6^S18/24A^ or Swi6^6S/A^ would have features that attract these mutant proteins. One possibility, especially at HOODs(Yamanaka et al. 2013) and islands (Zofall et al. 2012), is locally produced RNA(Keller et al. 2012), which may be more tightly bound by phospho-mutant versions of Swi6 (Stunnenberg et al. 2015; Nishibuchi et al. 2014). The general observation that loss of phosphorylation drives HP1 away from heterochromatin is consistent with data from human HP1α (Hiragami-Hamada et al. 2011) and *in vivo* diffusion measurements in the swi6 sm-1 mutant (Williams et al. 2024). This mutant likely disrupts NTE phosphorylation and shows greater residence outside heterochromatin. Further, it is likely that increased oligomerization of pSwi6 additionally strengthens this partitioning onto heterochromatin (next section). A separate consequence of this affinity decrease is the relief of competition with Clr4 for the nucleosome substrate (Figure 6, see third section below).

### Swi6 phosphorylation increases oligomerization

Swi6 has been shown to form dimers and higher-order oligomers. Swi6 oligomerization across chromatin has been linked to heterochromatin spreading *in vivo* (Canzio et al. 2011). Here, we show that phosphorylation increases the fraction of oligomeric states, revealing octamers and possibly higher molecular weight species (Figure 3). Swi6 exists in a closed dimer that inhibits the spreading competent state, or an open dimer that promotes oligomerization via the ARK loop in the CD (Canzio et al. 2013). One way pSwi6 could form higher molecular weight oligomers is by NTE phosphorylation shifting the equilibrium from the closed dimer to the open dimer, allowing the ARK loop access to engage a neighboring CD (Canzio et al. 2013) (Figure 7E). Our double mutant data (Figure 3E-J) supports such a model, whereby pS18 and/or pS24 cooperate with the ARK loop. We speculate the following thermodynamic consequence of phosphorylation on the nuclear Swi6 pool: Oligomerization will be driven at sites of high Swi6 accumulation, which is likely near its high-affinity H3K9me3 nucleosome target. If this is true, oligomerization will reduce the pool of free Swi6 available to engage unmethylated nucleosomes even further, and below the theoretical level we described above (∼5%).

HP1 proteins, like Swi6, form foci *in vivo*, which are associated with condensate formation, rooted in HP1 oligomerization (Larson et al. 2017; Strom et al. 2017; Sanulli et al. 2019). The reduction of GFP-Swi6S18/24A in nuclear foci we observe (Figure 3) may be due to defects in condensate formation, or simply that fewer Swi6 molecules are available to form heterochromatic condensates.

### Phosphorylation of Swi6 enables H3K9 trimethylation by Clr4

Achieving H3K9 trimethylation is essential for both gene silencing and heterochromatin spreading by Suv39/Clr4 enzymes (Muller et al. 2016; Al-Sady et al. 2013). For heterochromatin spreading, this is due to the positive feedback loop within Suv39/Clr4, which depends on binding trimethyl H3K9 tails via the CD (Al-Sady et al. 2013; Jih et al. 2017; Murawska et al. 2021).

Suv39/Clr4-mediated H3K9me0 to me1 and H3K9me1 to me2 conversion is 10X faster than the conversion from H3K9me2 to me3(Al-Sady et al. 2013). This slow step requires significant residence time on the nucleosome and is thus highly sensitive to factors promoting or antagonizing Clr4 substrate access, as well as nucleosome density (Cutter DiPiazza et al. 2021). Clr4 and Swi6 both make extensive contacts with nucleosomal DNA and the octamer core (Akoury et al. 2019; Sanulli et al. 2019). unpSwi6 and Clr4 affinity to H3K9me0 nucleosomes are very similar (1.8µM and 2.3µM for Clr4 (Al-Sady et al. 2013) and Swi6, respectively), but nuclear Swi6 concentration (2-4 µM) is likely higher than the Clr4 concentration (Iglesias et al. 2020). Thus, unpSwi6 would compete and displace Clr4 from its substrate. However, pSwi6’s affinity for the H3K9me0 nucleosome (28µM) is in a regime that is well above its predicted *in vivo* concentration. Any residual competition between pSwi6 and Clr4 would be mitigated by this lower affinity and the likely higher affinity of the Clr4 complex to its *in vivo* nucleosome substrate, driven by additional chromatin modfications (Stirpe et al. 2021).

This lowered pSwi6 nucleosome affinity likely relieves the trimethylation inhibition we observe for unpSwi6 (Figure 6, SFigure 9). Therefore, we propose that a major outcome of Swi6 phosphorylation is to clear nucleosome surfaces for Clr4 to access its substrate (Figure 7E). We note that instead of starting with H3K9me0 nucleosomes, we examined the conversion of H3K9me2 to me3. pSwi6 may have increased affinity to those H3K9me2 than H3K9me0 substrates (Sadaie et al. 2008). However, it has been shown by us (SFigure 4C) and others (Jih et al. 2017) that pSwi6 or Swi6 isolated from *S. pombe* cells, which is mostly phosphorylated (Figure 6A), has a preference for H3K9me3 over H3K9me2 (Jih et al. 2017). This lower H3K9me2 preference may still help pSwi6 distinguish between binding H3K9me2 versus me3 chromatin *in vivo*, and not just H3K9me0 versus H3K9me3.

Our *in vivo* data reveal several serines in Swi6 contribute to spreading, but S18 and S24 have a dominant effect. Why might these two residues, when mutated, have a strong impact on heterochromatin spreading? It is possible that pS18 and/or pS24 plays a disproportional role versus other residues in shifting the Swi6 from the closed to the open state, potentially blocking the ARK loop’s access to an adjacent CD when unphosphorylated. Alternatively, it is possible that *in vivo*, pS18 and/or pS24 are involved in the recruitment of H3K9me3-promoting factors, including Clr3, and also other factors like Abo1(Yamada et al. 2005; Zofall et al. 2022; Dong et al. 2020). Prior work (Shimada et al. 2009) has shown that Clr3 recruitment to heterochromatin is somewhat compromised in swi6S18-117A, while the recruitment of the anti-silencing protein Epe1 is increased. While this loss of Clr3 and gain of Epe1 may be an indirect consequence of compromised heterochromatin in swi6S18-117A, it cannot be excluded that phosphorylation at S18 and S24 is necessary to help recruit Clr3 and/or exclude Epe1. This would provide another mechanism for Swi6 to support trimethylation spreading by Clr4. Whether this is the case requires further investigation.

Together, we believe that our work resolves a critical problem in heterochromatin biology, which is how “writers” and “readers” promote heterochromatin spreading if they compete for the same substrate surfaces. Phosphorylation of Swi6 tunes the partitioning of Swi6 between unmethylated and methylated nucleosomes *in vivo*, such that Clr4 unmethylated substrates remain largely unbound. Whether this phosphorylation is regulated temporally, at different stages of heterochromatin formation, or spatially, at nucleation versus spreading sites, remains to be investigated.

## Material and Methods

### Strain construction

To construct wildtype swi6 and swi6 phosphoserine mutants, the swi6 open reading frame (ORF) was first deleted by integrating a ura4 gene cassette in the MAT HSS background. A plasmid, pRS316, was constructed containing 5’ homology-swi6 promoter-swi6 (or swi6 S-A mutant)-3’ UTR-kan-3’ genome homology and linearized by PmeI double digest to replace the ura4 cassette by genomic integration via homologous recombination. After transformation, cells were plated on YES agar for 24 hours before replica plating on G418 selection plates. For the □ckb1 mutant, we crossed the deletion strain from our chromatin function library(Greenstein et al. 2022) to the MAT □REIII□HSS and selected □ckb1 MAT □REIII□HSS strainsby random spore analysis on HYG+G418 double selection. For Swi6GFP fusions in the Sad1-mKO2 background, swi6 wildtype and swi6 S-A mutant strains were first crossed with the sad1::mkO2 strain to remove the MAT HSS. Next, swi6 and swi6 S-A mutant ORFs were C-terminally fused to SF-GFP followed by a hygromycin resistance marker by CRISPR/Cas9 editing as previously described (SpEDIT (Torres-Garcia et al. 2020)). Modifications were confirmed by gDNA extraction and PCR amplification of the 5’ swi6 to 3’ genome region downstream.

For the 3XFLAG tagged strains, a forward ultramer primer containing the swi6 5’UTR-3XFLAG-3’ swi6 ORF homology and a modified PAM sequence was used to amplify the 3XFLAG HR template to insert the N-terminal 3XFLAG tag into wildtype *swi6, swi6*^S18/24A^, and swi6^6S/A^ using CRISPR/Cas9 editing. The SpEDIT technology was also used to construct the loopX (*swi6*^R93A/K94A^) and acidicX (*swi6*^E(74-80)A^) mutations in wildtype swi6 and in combination with *swi6*^S18/24A^. For all strain construction, isolates were verified by genomic PCR.

### Western blot

Proteins were separated on a 15% SDS-Page gel and transferred to a PVDF membrane (Millipore) for 90 minutes at 100V and 4°C. Membranes were blocked overnight in 1:1 1X PBS: Intercept PBS Blocking Buffer (LiCor). Next, membranes were incubated with either polyclonal anti-Swi6 antibody (Canzio et al. 2013) or anti-pSwi6 antibody (Rockland Immunochemicals, this study) diluted 1:1000 in 1:1 1X PBS, 0.2% Tween-20 (PBS-T): Intercept PBS Blocking Buffer overnight at 4°C on a nutator. Anti-α-tubulin antibody was diluted 1:2000 and used as loading control. Membranes were washed twice with PBS-T for 10 minutes followed by two washes for 5 minutes before incubation with secondary antibodies. Secondary fluorescent antibodies were diluted either 1:10000 (anti-rabbit, 680 nm, Cell Signaling Technology 5366P, lot # 14) or 1:5000 (anti-mouse, 800 nm, Li-Cor, D10603-05) and were incubated with the membranes for 45 minutes at RT. Finally, membranes were washed 3 times with PBS-T for 10 minutes and once with PBS for 10 minutes before imaging on a LiCor Odyssey CLx imager.

### Flow cytometry

Strains were struck out of a -80°C freezer onto YES plates. Recovered cells were grown in 200 µL of YES media in a 96-well plate overnight to saturation at 32°C. The next morning, cells were diluted 1:25 in YES media into mid-log phase and analyzed by flow cytometry on an LSR Fortessa X50 (BD Biosciences). Fluorescence compensation, data analysis, and plotting in R were performed as described in Greenstein et al. 2022.For gene expression analysis in single cells, flow cytometry (FC) analysis was performed according to a previously described protocol (Greenstein et al. 2018).

### Chromatin immunoprecipitation followed by sequencing (ChIP -seq) sample collection and library preparation

For FLAG antibody (Figure 5 and SFigure 7) and pS18-pS24 antibody (SFigure 8) ChIP, strains were seeded in the morning, back diluted in the evening to OD600 ∼0.1 in 50mL YES media, and grown over night to saturation (32°C, 225RPM shaking). The following morning, cells at OD600 ∼1.8-2 were incubated at 18°C for 2 hours (225RPM shaking). 600×106 cells were pelleted and resuspended in 4.5 mL room temperature 1XPBS. To fix the cells, 1.5mM EGS (final) dissolved in DMSO was added to each sample and incubated at room temperature for 30 minutes followed by another 30min in 1% formaldehyde (final). Samples were quenched with 250mM glycine. Fixed cells were pelleted and washed twice with ice cold 1XTBS flash freezing and storing at -80°C.

For H3K9me2 antibody and H3K9me3 antibody ChIP, (data in Figure 2 and SFigure 2) cells were grown in YES media overnight to saturation (32°C, 225RPM shaking). The following morning cells were diluted to OD 0.03, grown to OD 1, and 300×106 cells were fixed only with formaldehyde, as above, and frozen at -80□C. Cells for FLAG, pS18-pS24, H3K9me2, and H3K9me3 ChIP were processed as described in Canzio et al. 2011 with the following modifications: Three technical replicates were processed for ChIPseq. For pS18-pS24 antibody ChIP, the lysis buffer was further supplemented with phosphatase inhibitors: 0.1 mM Sodium Orthovanadate and 60 mM β-Glycerophosphate disodium salt pentahydrate. After lysis, cells were bead beat 10 rounds for 1 minute each round with 0.5 mm Zirconia/Silica beads (Cat No. 11079105z). Tubes were chilled on ice for 2 minutes between rounds. Lysates were then spun down to isolate chromatin. The chromatin pellet was resuspended in 1.5 mL lysis buffer, moved to a 15 mL Diagenode Bioruptor tube (Cat. No. C01020031) and sonicated with a Diagenode Bioruptor Pico sonicator for a total of 35 cycles, 30 seconds on/ 30 seconds off, in the presence of sonication beads (Diagenode, Cat. No. C03070001). Every 10 cycles tubes were vortexed. Chromatin lysate was spun down for 30 minutes at 14000 RPM and 4°C. For H3K9me2 and H3K9me3 antibody ChIP, the lysate volume was brought up to 900 µL. 45 µL was taken out to check shearing of the DNA. 40 µL was taken out for input and kept at RT until the reverse crosslinking step. The remaining ∼800 µL was divided into 2 tubes to incubate with either 2 µL anti-H3K9me2 (Abcam 1120, Lot No. 1009758-6) or 1 µg anti-H3K9me3 (Diagenode, Cat. No. C15500003 Lot No. 003) overnight on a tube rotator at 4°C. For FLAG and pS18-pS24 anitbody ChIP, the lysate volume was brought up to ∼1.3mL. 45 µL was taken out to check shearing of the DNA. 40 µL was taken out for input and kept at RT until the reverse crosslinking step. The remaining ∼1.2mL was incubated with 2 µL M2 FLAG antibody (SIGMA cat: F1804, lot:0000308215) or 2 µL pS18-pS24 antibody, 1:1500 spike-in Drosophila DNA (1 ng spike-in DNA to 1500 ng input DNA quantified by Qubit) (Active Motif, Cat. No. 53083, Lot No. 24352138) and spike-in antibody (Active Motif, Cat. No. 61686, RRID: AB_2737370).

The next morning, Protein G Dynabeads (Invitrogen, LOT 3094426), Protein A Dynabeads (Invitrogen, LOT 01102248) and M-280 Streptavidin beads (Invitrogen, LOT 2692541) were washed twice with Lysis Buffer without protease inhibitors. 30 µL Protein G Dynabeads were added to each anti-FLAG and antipS18-pS24 sample, 20 µL Protein A Dynabeads beads were added to each anti-H3K9me2 sample, and 30 µL M-280 Streptavidin beads were added to each anti-H3K9me3 sample. Beads were incubated with samples for 3 hours on a tube rotator at 4°C, and then washed with 700 µL cold buffers at RT on a tube rotator in the following order: 2X Lysis Buffer for 5 minutes, 2X High Salt Buffer for 5 minutes, 1X Wash Buffer for 5 minutes, and 1X TE (buffer recipes as in [(Rougemaille et al. 2008)]). Samples were incubated with 100 µL elution buffer (50 mM Tris pH 8.0, 10 mM EDTA, 1% SDS) for 20 minutes at 70°C in a ThermoMixer F1.5 (Eppendorf). Input samples were brought up to 100 µL in TE with a final concentration of 1% SDS. Input and eluted samples were then incubated overnight in a 65°C water bath with 2.5 µL 2.5 mg/mL Proteinase K (Sigma Aldrich, Lot 58780500) for reverse crosslinking. Samples were purified with a PCR clean-up kit (Machery-Nagel) and eluted in 100 µL 10 mM Tris pH 8.0. The quality and size of the DNA were assessed by 4200 TapeStation instrument (Agilent). Next, libraries were prepared using NEBNext Multiplex Oligos for Illumina (E7335L, Lot 10172541; #E6442S, Lot #10236110; and #E6440S lot #10188082) and Ultra II FS DNA Library Prep Kit for Illumina (E7805L, Lot 10202083). The manufacturer’s protocol, “Protocol for FS DNA Library Prep Kit (E7805, E6177) with Inputs ≤100 ng (NEB)”, was used starting with 200 pg of DNA. PCR-enriched adaptor-ligated DNA was cleaned up using NEBNext sample purification beads (E6178S, Lot 10185312, “1.5. Cleanup of PCR Reaction” in manufacturer’s protocol). Individual adaptor-ligated DNA sample concentrations were quantified using a Qubit 4 Fluorometer (Thermo Fisher), and the quality of the DNA was assessed by a 4200 TapeStation instrument (Agilent). Libraries were pooled to equimolar quantities and sequenced using either a NextSeq 2000 P2 (400 million clusters) (Chan Zuckerberg Biohub San Francisco) (40bp read length, paired-end) for Histone ChIP-seq, or a Novaseq X plus (25B 300 chemistry) (Signios) (150bp read length, paired-end) for Flag-Swi6 and Phospho-Swi6.

### ChIP-seq data analysis

Sequencing adaptors were trimmed from raw sequencing reads using Trimmomatic v0.39.

The *S. pombe* genome was downloaded from NCBI under Genome Assembly ASM294v2. The MAT locus of chromosome II was edited to our custom HSS MAT locus, and the genome was indexed using the bowtie2-build function of Bowtie2 v2.5.1(Bolger et al. 2014). For H3K9me2/3 ChIP, trimmed sequencing reads were aligned to the genome using Bowtie2 v2.5.1 with flags [--local -- very-sensitive-local --no-unal --no-mixed --no-discordant -- phred33 -I 10 -X 700](Langmead and Salzberg 2012). Next, the resulting SAM files were converted to BAM files using SAMtools v1.18(Li et al. 2009) view function: -S -b ${base}.sam > ${base}.bam. The resulting BAM files were further processed by removing low-quality alignments, PCR duplicates, and multimappers, and retain properly aligned paired-end reads using SAMtools view with the following flags: -bh -F 3844 -f 3 -q 10 -@ 4. The processed BAM files were then sorted and indexed (SAMtools). Sorted, indexed BAM files were converted to bigWig coverage tracks using deepTools v3.5.4(Ramírez et al. 2014): bamCoverage: -b “$bam_file” -o “$filename_without_extension.bw” --binSize 10 --normalizeUsing CPM --extendReads --exactScaling --samFlagInclude 64 --effectiveGenomeSize 13000000. BigWig files normalized to input were generated using the bigwigCompare tool (deepTools). For FLAG-Swi6 and pS18-pS24 Swi6 ChIP, trimmed reads were mapped using Bowtie2 v2.5.3 using default parameters. Output SAM files were converted to BAM files, sorted, and indexed using samtools v1.21 sort, view, and index functions. Spike in was mapped using bowtie2 v2.5.3 with the BDGP6 bowtie2 index for D. melanogaster using default values. Spike-in normalization was performed by taking the “aligned concordantly exactly 1 time” read count and dividing by the lowest value. Coverage tracks were generated as above, but removed the SamFlagInclude option and adding the scaleFactor option with the calculated scale factor for each library. Normalized bigWig files were loaded into R v4.3.0 using rtracklayer v1.60.1(Lawrence et al. 2009) and processed for visualization as in Greenstein et al. 2022 with modifications. The Gviz v1.44.2 (Bioconductor) DataTrack function was used to create a visualization track of ChIP-seq signal in bigWig files for each genotype(Hahne and Ivanek 2016). The Bioconductor GenomicRanges package was used to create a GRanges object to store custom genomic coordinates defined by a BEDfile([CSL STYLE ERROR: reference with no printed form.]). Genomic annotations for signal tracks were created using the AnnotationTrack (Gviz) function. The GenomeAxisTrack (Gviz) function generated a visual reference (in megabases) to display the position of genomic annotations and signal tracks. Finally, the plotTracks (Gviz) function was used to plot the DataTrack, AnnotationTrack, and GenomeAxisTrack objects for visualization. Underlying values for the heatmaps were calculated by using BigWigAverageoverBed of spike-in normalized signal tracks and coordinate files for each annotated region. Output was visualized using GraphPad Prism

#### Swi6 phosphorylation enables spreading

v10.6. For differential enrichment, reads were counted into 300bp windows by the Bioconductor package csaw and background normalized using 5kb windows and the normFactors() command. TMM count normalization and differential enrichment were performed by the edgeR function. Output was visualized using ggplot2. Annotations for regions were generated and intersected as described in Greenstein et al. 2022.

### Swi6-GFP live cell imaging

Swi6-GFP/Sad1-mKO2 strains were struck out onto fresh YES 225 agar plates and incubated at 32°C for 3-5 days. Colonies were inoculated into liquid YES 225 medium (#2011, Sunrise Science Production) and grown in an incubator shaker at 30°C, 250 rpm to an OD of 0.2 -0.6. Cells were placed onto 2% agarose (#16500500, Invitrogen) pads in YES 225, covered with a coverslip (#2850-22, thickness 1 ½, Corning), and sealed with VALAP for imaging. Cells were imaged on a TiEclipse inverted microscope (Nikon Instruments) with a spinning-disk confocal system (Yokogawa CSU-10) and a Borealis illumination system that includes 488nm and 541nm laser illumination and emission filters 525±25nm and 600±25nm respectively, 60X (NA: 1.4) objectives, and an EM-CCD camera (Hamamatsu, C9100-13). These components were controlled with μManager v. 1.41(Edelstein et al. 2014, 2010). The temperature of the sample was maintained at 30°C by a black panel cage incubation system (#748–3040, OkoLab). The middle plane of cells was first imaged in brightfield and then two Z-stacks with a step size of 0.5µm were acquired in spinning-disk confocal mode with laser illumination 488nm and 541nm (total of 9 imaging planes per channel). The exposure, laser power, and EM gain for the Z-stacks were respectively 50ms / 1% / 800, and 200ms / 5% / 800. Between 9 and 12 fields of view were acquired per strain.

### Image analysis

For each field of view, nuclei were manually cropped using Fiji. Cells containing multiple Sad1-mK02 foci were discarded from this analysis. For each selected nucleus TrackMate was used to determine the coordinates in space (X, Y, Z) of Sad1 and every Swi6 focus and their fluorescence intensity(Ershov et al. 2022; Tinevez et al. 2017). Using a custom script on Jupiter Notebook in Python we then automatically counted the number of Swi6 foci detected by TrackMate for each nucleus. Additionally, we automated the calculation of the distance between Swi6 foci and the spindle pole body by measuring the distance from each Swi6 focus to Sad1 within a given nucleus. For Swi6 intensity measurements, a region of interest (ROI) outside of each nucleus was automatically selected to measure the background intensity. This background intensity was then used to correct Swi6 fluorescence signal by subtracting the average intensity of this ROI for a given analyzed nucleus. Finally, we used a one-way ANOVA statistical test on Swi6 intensity signal to determine differences between mutants.

### Protein cloning and purification

Wildtype swi6 open reading frame was cloned by ligation-independent cloning into vector 14B (QB3 Berkeley Macrolab expression vectors). Vector 14B encodes an N-terminal 6xHis tag followed by a TEV cleavage sequence. Wildtype Swi6 was expressed in BL21-gold (DE3) competent cells. To produce Swi6S18/24A, a gene block containing S18A/S24A Swi6 was cloned into vector 14B using Gibson assembly. To isolate pSwi6 and pSwi6S18/S24A, the respective vectors were co-expressed with the catalytic subunits of Caesin Kinase II in pRSFDuet. All three proteins were grown, harvested, and purified using a protocol adapted from [10] and modified as follows: Cells were grown at 37°C until OD600 0.5-0.6 and induced with a final concentration of 0.4mM Isopropyl-β-D-thiogalactopyranoside. Induced cells were grown at 18°C overnight. Harvested cells were resuspended in lysis buffer containing 1X PBS buffer pH 7.3, 300 mM NaCl, 10% glycerol, 0.1% Igepal CA-630, 7.5 mM Imidazole, 1 mM Beta-Mercaptoethanol (βME), with protease inhibitors. Resuspended cells were sonicated 2 seconds on /2 seconds off at 40% output power for three 5-minute cycles. The lysate was centrifuged at 25,000xg for 25 minutes, and the supernatant was collected. Nickel NTA resin was equilibrated with lysis buffer. The supernatant and resin were incubated for 1-2 hours and washed 3 times with 40 ml of lysis buffer each time before the protein was eluted with 25 mM HEPES pH 7.5, 100 mM KCl, 10% glycerol, 400 mM Imidazole, and 1 mM βME. The eluted protein was then dialyzed in TEV cleavage buffer containing 25 mM HEPES pH 7.5, 100 mM KCl, and 1 mM βME and 6 mg TEV protease. The following morning 3-6 mg of TEV protease was spiked in for about 1 hour to ensure full cleavage. Nickel NTA resin was equilibrated with TEV cleavage buffer and the his-tagged TEV was captured by the resin while Swi6 protein was isolated by gravity flow. Cleaved protein was concentrated using a 10kDa MWCO concentrator and applied to a Superdex 200 Increase 10/300 GL size exclusion column equilibrated in storage buffer containing 25 mM HEPES pH 7.5, 100 mM KCl, 10% glycerol, and 10 mM βME. Protein was concentrated, flash-frozen in N2 (liq), and stored at -80°C. Protein concentration was quantified against a BSA standard curve on an SDS page gel and sypro red stain.

### Crosslinking

SEC-MALS: unpSwi6 or pSwi6 was purified as described above. However, the storage buffer was 25 mM HEPES pH 7.5 and 100 mM KCl for SEC-MALS. Protein, either 100 µM Swi6 or pSwi6, was incubated with 2 mM EDC and 5 mM NHS in a total volume of 95 µL for 2 hours. The reaction was quenched with a final concentration of 20 mM hydroxylamine.

Crosslinking western: To compare oligomerization states in pSwi6 and pSwi6S18/24A, the protein was purified as above into storage buffer and crosslinked at various concentrations for 2 hours at room temperature in 0.01% glutaraldehyde. Crosslinking was quenched with 50mM Tris pH 7.4 and the protein diluted to 4.2µM before separation on a 4-12% Tris-glycine gradient gel (Novex, Invitrogen). Western was performed as above.

### Size-exclusion Chromatography coupled with Multi-Angle Light Scattering (SEC-MALS)

Crosslinked and uncrosslinked Swi6 and pSwi6 were filtered with 0.2 µm spin columns (Pall Corporation, Ref. ODM02C34). For SEC, uncrosslinked and crosslinked proteins were injected onto a KW-804 silica gel chromatography column (Shodex) in a volume of 50 µL at 100 µM. The column was run using an ÄKTA pure FPLC (GE Healthcare Life Sciences) and equilibrated with SEC-MALS storage buffer at a flow rate of 0.4 mL/min and temperature of 8°C. The SEC column was connected in-line to a DAWN HELEOS II (Wyatt Technology) 18-angle light scattering instrument and an Optilab T-rEX differential refractive index detector (Wyatt Technology). Data was analyzed using ASTRA software (version 7.1.4.8, Wyatt Technology) and graphed using GraphPad Prism software (version 9.5.1).

### Fluorescence Polarization

Fluorescence Polarization binding measurements were conducted as described in Canzio et al. 2013 and modified as follows: Peptide reaction buffer was 50 mM HEPES pH 7.5, 100 mM KCl, 10% glycerol, 0.01% NP-40, and 2 mM βME. Fluoresceinated H31–20 K9me0 or H31–20 K9me3 peptide concentration was fixed at 100 nM while Swi6, pSwi6, or pSwi6S18/24A protein concentration varied from 0-200 µM. Mononuclesome reaction buffer was 20 mM HEPES pH 7.5, 80 mM KCl, 4 mM Tris, 0.2 mM EDTA, 10% glycerol, 0.01% NP-40, 2 mM βME. H3K9me0 and H3KC9me3 mononucleosomes were reconstituted with fluorescein-labeled 601 DNA as described (Canzio et al. 2011). Nucleosome concentration was fixed at 25 nM while Swi6, pSwi6, or pSwi6S18/24A protein concentrations varied from 0-200 µM. Both peptide and mononucleosome reaction volumes were 10 µL and measured in a Corning 384 low-volume, flat bottom plates (product number 3820, LOT 23319016). Fluorescence polarization was recorded using a Cytation 5 microplate reader (Biotek, λex= 485/20nm, λem= 528/20nm) and Gen5 software (Biotek, version 3.09.07). Data was analyzed and fit to a Kd equation using GraphPad Prism.

### Single turnover kinetics

Clr4 protein was purified exactly as described (Al-Sady et al. 2013). H3K9me2 nucleosomes were purchased from Epicypher (#16-0324), and pSwi6 and unpSwi6 were purified as above. Single turnover reactions were carried out as follows: 5 µM Clr4 was preincubated 5 minutes with 1 mM final S-adenosyl-methionine (liquid SAM, 3 2mM, NEB #B9003S), and varying concentrations of pSwi6, pSwi6S18/24A, or unpSwi6, at 25°C to reach equilibrium. 5µM Clr4 was chosen as the minimal Clr4 concentration to yield robust H3K9me3 signal under Single Turnover conditions. The reaction was started with the addition of H3K9me2 nucleosomes to 500nM final. Timepoints were stopped by boiling with SDS-Laemmli buffer. Samples were separated on 18% SDS-PAGE gel and probed for the presence of H3K9me3 (polyclonal, Active Motif #39161. lot 22355218-11) and H4 (Active Motif #39269 lot 31519002) as a loading control. Signals were quantified on a Li-Cor imager by using a dilution of H3K9me3 nucleosomes (Epicypher, #16-0315), establishing standard curves for H4 and H3K9me3. Rates were fit to a single exponential rise in GraphPad Prism software exactly as published (Al-Sady et al. 2013).

### Phosphatase treatments

1500 ng of Swi6, pSwi6, and pSwi6^S18/24A^ were incubated for 20 minutes at 37°C with 50 U of Calf Intestinal Phosphatase (QuickCIP, NEB, M0525S) in 100 mM NaCl, 50 mM Tris-HCl, 10 mM MgCl2, 1 mM dithiothreitol, pH 7.9. Reactions were stopped by boiling in SDS-Laemmli buffer. For reactions with inactivated CIP, 200U CIP was pre-incubated for 20 minutes at 80°C. 75 ng of Swi6, pSwi6, and pSwi6^S18/24A^ that was either mock-treated, treated with active or inactivated CIP was separated on either a 15% SDS-PAGE gel or a SuperSep Phos-Tag gel (Fujifilm, 15.5%, 17 well, 100×100×6.6mm, Lot PAR5302). The Phos-Tag gel was washed with western transfer buffer with 10 mM EDTA to remove Zn2+ ions and then blotted and probed for Swi6 with Swi6 polyclonal antibody as above.

### Mass Spectrometry

In-solution Trypsin/Lys C digested peptides were analyzed by online capillary nanoLC-MS/MS using several different methods. High resolution 1 dimensional LCMS was performed using a 25 cm reversed-phase column fabricated in-house (75 µm inner diameter, packed with ReproSil-Gold C18-1.9 μm resin (Dr. Maisch GmbH)) that was equipped with a laser-pulled nanoelectrospray emitter tip. Peptides were eluted at a flow rate of 300 nL/min using a linear gradient of 2–40% buffer B in 140 min (buffer A: 0.02% HFBA and 5% acetonitrile in water; buffer B: 0.02% HFBA and 80% acetonitrile in water) in a Thermo Fisher Easy-nLC1200 nanoLC system. Peptides were ionized using a FLEX ion source (Thermo Fisher) using electrospray ionization into a Fusion Lumos Tribrid Orbitrap Mass Spectrometer (Thermo Fisher Scientific). Data was acquired in orbi-trap mode. Instrument method parameters were as follows: MS1 resolution, 120,000 at 200 m/z; scan range, 350−1600 m/z. The top 20 most abundant ions were subjected to higher-energy collisional dissociation (HCD) or electron transfer dissociation (ETD) with a normalized collision energy of 35%, activation q 0.25, and precursor isolation width 2 m/z. Dynamic exclusion was enabled with a repeat count of 1, a repeat duration of 30 seconds, and an exclusion duration of 20 seconds.

Low-resolution, 1-dimensional LCMS was performed using a nano-LC column packed in a 100-μm inner diameter glass capillary with an integrated pulled emitter tip. The column consisted of 10 cm of ReproSil-Gold C18-3 μm resin (Dr. Maisch GmbH)). The column was loaded and conditioned using a pressure bomb. The column was then coupled to an electrospray ionization source mounted on a Thermo-Fisher LTQ XL linear ion trap mass spectrometer. An Agilent 1200 HPLC equipped with a split line so as to deliver a flow rate of 1 ul/min was used for chromatography. Peptides were eluted with a 90-minute gradient from 100% buffer A to 60% buffer B. Buffer A was 5% acetonitrile/0.02% heptafluorobutyric acid (HBFA); buffer B was 80% acetonitrile/0.02% HBFA. Collision-induced dissociation spectra were collected for each m/z.

Multidimensional protein identification technique (MudPIT) was performed as described(Liu et al. 2002; Washburn et al. 2001). Briefly, a 2D nano-LC column was packed in a 100-μm inner diameter glass capillary with an integrated pulled emitter tip. The column consisted of 10 cm of ReproSil-Gold C18-3 μm resin (Dr. Maisch GmbH)) and 4 cm strong cation exchange resin (Partisphere, Hi Chrom). The column was loaded and conditioned using a pressure bomb. The column was then coupled to an electrospray ionization source mounted on a Thermo-Fisher LTQ XL linear ion trap mass spectrometer. An Agilent 1200 HPLC equipped with a split line so as to deliver a flow rate of 1 ul/min was used for chromatography. Peptides were eluted using a 4-step gradient with 4 buffers. Buffer (A) 5% acetonitrile, 0.02% heptafluorobutyric acid (HFBA), buffer (B) 80% acetonitrile, 0.02% HFBA, buffer (C) 250mM NH4AcOH, 0.02% HFBA, (D) 500mM NH4AcOH, 0.02% HFBA. Step 1: 0-80% (B) in 70 min, step 2: 0-50% (C) in 5 min and 0-45% (B) in 100 min, step 3: 0-100% (C) in 5 min and 0-45% (B) in 100 min, step 4 0-100% (D) in 5 min and 0-45% (B) in 160 min. Collision-induced dissociation (CID) spectra were collected for each m/z.

Data analysis: RAW files were analyzed using PEAKS (Bioinformatics Solution Inc) with the following parameters: semi-specific cleavage specificity at the C-terminal site of R and K, allowing for 5 missed cleavages, precursor mass tolerance of 15 ppm (3 Da for low-resolution LCMS), and fragment ion mass tolerance of 0.5 Daltons. Methionine oxidation and phosphorylation of serine, threonine, and tyrosine were set as variable modifications and Cysteine carbamidomethylation was set as a fixed modification. Peptide hits were filtered using a 1% false discovery rate (FDR). Phosphorylation occupancy ratio for amino acids was determined by summing the count of unphosphorylated and phosphorylated amino acids detected in the experiment. We only considered phospho-peptides detected more than once and at least 2% minimal ion intensity.

To assess the percent of peptides phosphorylated at S18 and/or S24 more precisely, we performed three separate Mass Spectrometry experiments with the pSwi6 protein from independent purifications. Using PEAKS software, we counted all instances of peptides containing S18 and/or S24 phosphorylation, over peptides spanning the region without S18 or S24 phosphorylation. We did so in either Analysis mode 1 (semi-specific digest mode, which considers peptides with at least one tryptic cleavage end) or mode 2 (specific digest mode, which considers only peptides with two tryptic cleavage ends). Because our pS18-pS24 antibody is raised against an pS18-pS24 peptide, we cannot distinguish whether it recognizes S18 and/or S24 phosphorylation; hence, we report pS18 and/or pS24.

### Estimate of *in vivo* nucleosome fractions bound

*In vitro*binding isotherms for nucleosomes (N) can be fit simply via fraction Nbound=[Swi6]/([Swi6]+Kd). However, this assumes first that [N] << [Swi6] and << Kd, such that [Swi6] total [Swi6] free. In the nucleus, these assumptions do not hold. However, a quadratic equation (Jarmoskaite et al. 2020) can be used to estimate N bound, accounting for bound Swi6. In this case, frNbound=([N]total+[Swi6]total+Kd-√(?([N]total+[Swi6]total+Kd)?^2-4*[N]total*[Swi6]total))/(2*[N]total). To estimate the fraction of unmethylated nucleosomes bound by pSwi6 or unpSwi6, we used Kds from Figure 4F, total Swi6 concentrations of 2.1-4.6µM, and total nucleosome contraction estimate of ∼10µM.

## Supporting information

All Supporting material

## Data availability statement

ChIP-Seq data is deposited at NIH GEO Record GSE271394.

## Acknowledgements

We would like to thank Geeta J. Narlikar for the discussion on theory and critical feedback on the manuscript. We thank Sigurd Braun, Daniele Canzio, and Lucy D. Brennan for valuable feedback and discussion. B.A-S. was supported by a National Institutes of Health grant R35GM141888 and a National Science Foundation grant 2113319. D.R.K. and C.T. were supported by a National Science Foundation Graduate Research Fellowships Grant No. 2034836. E.S. was supported by a grant from the Ford Foundation and National Institutes of Health supplement DP2GM123484-01S1. E.M. was supported by an NIH IRACDA grant 5K12GM081266-17 and NIH T32 training grant 5T32HL007731-32. We acknowledge the UCSF PFCC (RRID:SCR_018206) for assistance in generating Flow Cytometry data. The research reported here was supported in part by the DRC Center Grant NIH P30 DK063720. We thank the Southworth lab for access and training to SEC-MALS equipment.

