## Supplementary material for "Phosphorylation of HP1/Swi6 relieves competition with Suv39/Clr4 on nucleosomes and enables H3K9 trimethyl spreading": All Supporting material

**Table 1.** Table of *S. pombe* strains used in this work.

**Supporting Figure 1: Additional isolates demonstrating that S18 and S24 in Swi6 are required for spreading, but not nucleation of heterochromatin silencing.** 2-D Density hexbin plots examining silencing at nucleation 'green' and spreading 'orange' reporter in the MAT *ΔREIII* HSS for three additional isolates of **A.-C.** $swi6^{S18/24A}$ mutants, **D.-F.** $swi6^{S46/5/117\text{-}220A}$ ("S18/24 available"), and **G.-I.** $swi6^{S46/52A}$. **J-L.** As A.-I. but for the *Δckb1* mutant. An independent wildtype isolate from the cross is shown alongside 2 *Δckb1* isolates. 'green' ON threshold based on Figure 1D-I.

**Supporting Figure 2: Heatmaps of H3K9me2 and H3K9me3 ChIP-seq. A.** heatmaps for H3K9me2 signal (IP over input; IP/Inp) at nucleators (top) and spreading zones (bottom) as previously defined[36]. **B.** as in A. but for H3K9me3.

**Supporting Figure 3: H3K9me2 and H3K9me3 ChIP-Seq plots in additional genomic loci in wildtype or** $swi6^{S18/24A}$. H3K9me2 (TOP) and H3K9me3 (BOTTOM) plots for wildtype and $swi6^{S18/24A}$ as in Figure 2 at **A.** *mei4,* **B.** *tel IL, and* **C.** *tel IIL*.

**Supporting Figure 4: Characterization of recombinant pSwi6 A.** Triplicate analysis of S18 and/or S24 phosphopeptide abundance using Mass Spectrometry. Analysis 1 = semi-specific digest mode; Analysis 2= specific digest mode. **B.** Size Exclusion Chromatography followed by Multi-Angle Light Scattering (SEC-MALS) on uncrosslinked unpSwi6 (black) and pSwi6 (green). Relative refractive index signals (solid lines, left y-axis) and derived molar masses (lines over particular species, right y-axis) are shown as a function of the elution volume. A migration shift is apparent in pSwi6, as well as a small shoulder of higher molecular weight species (arrow). **C.** FP of H3K9me2 (open circles) and H3K9me3 (filled circles) tail peptides with pSwi6. Derived affinities on the right.

**Supporting Figure 5: Additional isolates demonstrating that S18/S24 phosphorylation is working through, or in parallel to, known Swi6 oligomerization surfaces.** 2-D Density hexbin plots examining silencing at nucleation 'green' and spreading 'orange' reporter in the MAT *ΔREIII* HSS for two additional isolates of **A.** $swi6^{S18/24A}$ mutants (same as in SFigure 1 but run here in a separate experiment) **B.** $swi6^{loopX}$ (R93A K94A), **C.** $swi6^{loopX;S18/24A}$, **D.** $swi6^{acidicX}$ (E74-80A), and **E.** $swi6^{acidicX;S18/24A}$.

**Supporting Figure 6: Analysis of Swi6-GFP heterochromatin foci number and spatial distribution. A.** Strategy for production of GFP-tagged *swi6* S-A mutants in the *sad1:mKO2* background. The wildtype *swi6* or S-A mutant gene from Figure 2A was cut with CRISPR/Cas9, and the break was repaired with a cassette containing a super-folder GFP, *swi6* 3' sequence homology, and a HygMX cassette. **B.** Representative maximum projection live microscopy images of indicated Swi6$^{S46/52A}$-GFP/ Sad1-mKO2 compared to the wildtype strain. **C.** Quantification of Swi6-GFP signals by flow cytometry. The GFP signal of independent wildtype or S-A mutant isolates compared to GFP negative (GFP -) cells as measured by flow cytometry. **D.** Distribution of nuclear foci in nuclei of indicated strains represented as relative frequency. Wildtype Swi6-GFP , n=85; Swi6$^{S18/24A}$-GFP, n=94; Swi6$^{S18/24/117-220A}$-GFP, n=50; Swi6$^{S46/52/117-220A}$-GFP, n=82. **E.** distribution of Swi6-GFP heterochromatin foci relative to Sad1-mKO2. overview: center-to-center distances were measured in 3D from the peri-spindle pole body Sad1-mKO2 signal to all Swi6-GFP foci identified in each nucleus. **F.** relative frequency histogram binning the distribution of Sad1-mKO2 to Swi6-GFP foci distances in indicated strains.

**Supporting Figure 7: FLAG-Swi6 ChIP-Seq plots in additional genomic loci in FLAG-Swi6, FLAG-Swi6$^{S18/24A}$, and FLAG-Swi6$^{6S/A}$ strains**. FLAG ChIP Seq signal visualization plots as in Figure 5. The solid ChIP/input line for each genotype represents the mean of three repeats, while the shading represents the 95% confidence interval. **A.** *cen I* with zoom in of the left and right pericentromeric region. The brown dashed boxes indicate siRNAi-generating centers as mapped in [42]. **B.** As in A., for *tel IIR* and *tel IIL.* **C.** As in Figure 5G, except for all three genotypes and all replicates included.

**Supporting Figure 8: Swi6 pS18-pS24 ChIP-seq. A.** Schematic of the ChIP experiment using anti-pS18-pS24 antibody: Spike-in controlled ChIP-Seq was performed in wildtype *swi6* or swi6$^{S18/24A}$ mutant as a negative control for the specificity of the antibody. **B.** Top, Middle: Heatmaps of spike-in normalized pS18-pS24-ChIP-Seq signal (in Counts Per Million, CPM) for *swi6* and *swi6$^{S18/24A}$* for regions previously classified as nucleators or spreading zones[36] as in Figure 5B. Bottom: Euchromatic regions previously identified to constitutively or facultatively accumulate H3K9me2 (HOODs[59], islands[60], regions that accumulate H3K9me2 in *Δmst2Δepe1*[61]) irrespective of H3K9me2 status in our dataset. Note all features of *cenH*-inserted 'green' and MAT locus 'orange' are combined **C.- E.** Signal track visualization plots for FLAG-Swi6 (from Figure 5, right y-axis) and pS18-pS24 signals for *swi6* and *swi6$^{S18/24A}$* (left y-axis). **C.** ChIP-seq signal visualization plot for the MAT *ΔREIII* HSS mating type locus. The solid ChIP/input line for each genotype represents the mean of two repeats, while the shading represents the 95% confidence interval. **D.** As in C. but for *tel IIR.* **E.** As in C. but for *cen II* with zoom-in of the left and right pericentromeric region. The brown dashed boxes indicate siRNAi-generating centers as mapped in [42]. **F.** Volcano plots representing -log10(FDR) vs Log2 Fold Change for FLAG-Swi6 over pS18-pS24 Swi6 in wildtype. Dots represent 300bp windows tested for differential enrichment by edgeR. Colors indicate overlap of windows to annotated regions: HOODs, Islands, Regions; nucleators; spreading; or other. Cutoff values for FDR <0.01 and Log2 Fold Change of 1 are annotated on the plots. Right plot is the zoomed in region annotated on the left plot.

**Supporting Figure 9: Additional replicates of Swi6 westerns from cell lysates and nucleosome trimethylation assay. A.** Independent repeat of α-Swi6 and α-pS18-pS24 westerns as in Figure 4A. **B.** A repeat of quantitative western blots querying time-dependent formation of H3K9me3 from H3K9me2 mononucleosomes in the presence of pSwi6 or unpSwi6 (0 and 5μM Swi6). **C.** A repeat of quantitative western blots querying time-dependent formation of H3K9me3 from H3K9me2 mononucleosomes in the presence of pSwi6 or unpSwi6 (15μM and 30μM Swi6). **B**. and **C.** Swi6 concentration time courses were collected at the same time; westerns were run on separate days.

**Supporting Figure 10: pSwi6$^{S18/24A}$ mutant characterization. A.** Calf Intestine Phosphatase (CIP) treatment of Swi6 examined in a 15% SDS-PAGE gel. unpSwi6, pSwi6, or pSwi6$^{S18/24A}$ were treated with (+) or without (-) CIP or with heat-inactivated CIP (b). **B.** CIP treatment of Swi6 examined in a Phos-Tag gel as in A. Blots of both gels were probed with an anti-Swi6 polyclonal antibody. **C.** Mass Spectrometry on pSwi6$^{S18/24A}$. Shown is a domain diagram of Swi6$^{S18/24A}$. Phosphorylation sites identified in pSwi6 $^{S18/24A}$ by 2D-ETD-MS are indicated and grouped by detection prevalence in the sample. **D.** Derived affinities for nucleosome FP experiment in Figure 7C.

**Supporting Table 1**

Yeast strains used in this study

| Identifier | Genotype | Figure; experiment | Source |
|---|---|---|---|
| PAS210 | h+, *sad1*:mKO2:NATMX | Fig. 3G, H; SFig. 4 | Al-Sady *et al.* 2016 |
| PAS807 | h90, cenH:: ade6p:SF-GFP (Kint2); mat3m(EcoRV):: ade6p:mKO2; ade6p:3xE2C:hygMX at Locus2; ΔREIII::REIII(Δs1, Δs2), *swi6::ura4* | Fig. 1C,D; Fig. 2B; SFig. 2A,B; SFig. 3A-C | This study |
| PAS814 | h90, cenH:: ade6p:SF-GFP (Kint2); mat3m(EcoRV):: ade6p:mKO2; ade6p:3xE2C:hygMX at Locus2; ΔREIII::REIII(Δs1, Δs2), *swi6*:KANMX | Fig. 1C,E; Fig. 2B-D; Fig. 3E; SFig. 2A,B; SFig. 3A-C, SFig. 8B-F | This study |
| PAS851,858, 859, 860 (isolates) | h90, cenH:: ade6p:SF-GFP (Kint2); mat3m(EcoRV):: ade6p:mKO2; ade6p:3xE2C:hygMX at Locus2; ΔREIII::REIII(Δs1, Δs2), *swi6S18/24A*:KANMX | Fig. 1C,G; SFig.1A-C; Fig. 2B-D; SFig. 2A,B SFig.3 A-C, SFig. 8B-E | This study |
| PAS852, 861,862, 863 isolates) | h90, cenH:: ade6p:SF-GFP (Kint2); mat3m(EcoRV):: ade6p:mKO2; ade6p:3xE2C:hygMX at Locus2; ΔREIII::REIII(Δs1, Δs2), *swi6S46/52,117-220A*:KANMX | Fig. 1C,H; SFig.1D-F | This study |
| PAS853, 864, 865, 866 (isolates) | h90, cenH:: ade6p:SF-GFP (Kint2); mat3m(EcoRV):: ade6p:mKO2; ade6p:3xE2C:hygMX at Locus2; ΔREIII::REIII(Δs1, Δs2), *swi6S46/52*:KANMX | Fig. 1C,F | This study |
| PAS854 | h90, cenH:: ade6p:SF-GFP (Kint2); mat3m(EcoRV):: ade6p:mKO2; ade6p:3xE2C:hygMX at Locus2; ΔREIII::REIII(Δs1, Δs2), *swi6S18/24,117-220A*:KANMX | Fig. 1C,I | This study |
| PAS909 | *sad1*:mKO2:NATMX; *swi6S46/52*:SF-GFP:HYGMX | SFig. 4B-F | This study |
| PAS910,911 | *sad1*:mKO2:NATMX; *swi6S18/24,117-220A*:SF-GFP:HYGMX | Fig. 3G,H; SFig. 4C-F | This study |
| PAS913 | *sad1*:mKO2:NATMX; *swi6*:SF-GFP:HYGMX | Fig. 3G,H; SFig. 4C-F | This study |
| PAS919 | *sad1*:mKO2:NATMX; *swi6S18/24*:SF-GFP:HYGMX | Fig. 3G,H; SFig. 4C-F | This study |
| PAS922 | *sad1*:mKO2:NATMX; *swi6S46/52,117-220A*:SF-GFP:HYGMX | Fig. 3G,H; SFig. 4C-F | This study |
| PAS1189 | h90, cenH:: ade6p:SF-GFP (Kint2); mat3m(EcoRV):: ade6p:mKO2; ade6p:3xE2C:hygMX at Locus2; ΔREIII::REIII(Δs1, Δs2), Δ*ckb1::*KANMX | Fig. S1K,L | This study |
| PAS1195, PAS1196, PAS1197 (isolates) | h90, cenH:: ade6p:SF-GFP (Kint2); mat3m(EcoRV):: ade6p:mKO2; ade6p:3xE2C:hygMX at Locus2; ΔREIII::REIII(Δs1, Δs2), 3XFLAG-*swi6*:KANMX | Fig. 5B-G, SFig. 3A-C, SFig. 7A-C, SFig. 8C-F | This study |
| PAS1198, PAS1199, PAS1200 (isolates) | h90, cenH:: ade6p:SF-GFP (Kint2); mat3m(EcoRV):: ade6p:mKO2; ade6p:3xE2C:hygMX at Locus2; ΔREIII::REIII(Δs1, Δs2), 3XFLAG-*swi6S18/24A*:KANMX | Fig. 5B-G; SFig. 3A-C; SFig. 7A-C; SFig. 8B-F | This study |
| PAS1202, PAS1203, PAS1204 (isolates) | h90, cenH:: ade6p:SF-GFP (Kint2); mat3m(EcoRV):: ade6p:mKO2; ade6p:3xE2C:hygMX at Locus2; ΔREIII::REIII(Δs1, Δs2), 3XFLAG-*swi6S18/24,117-220A*:KANMX | Fig. 5B-F; SFig. 7A-C | This study |
| PAS1206 (isolate1, 2, 3) | h90, cenH:: ade6p:SF-GFP (Kint2); mat3m(EcoRV):: ade6p:mKO2; ade6p:3xE2C:hygMX at Locus2; ΔREIII::REIII(Δs1, Δs2), *swi6R93, K94A*:KANMX | Fig. 3G,;SFig5B | This study |
| PAS1207 (isolate 1, 2, 3) | h90, cenH:: ade6p:SF-GFP (Kint2); mat3m(EcoRV):: ade6p:mKO2; ade6p:3xE2C:hygMX at Locus2; ΔREIII::REIII(Δs1, Δs2), *swi6S18/24A,R93A,K94A*:KANMX | Fig. 3H; SFig5C | This Study |
| PAS1210 (isolate 1, 2, 3) | h90, cenH:: ade6p:SF-GFP (Kint2); mat3m(EcoRV):: ade6p:mKO2; ade6p:3xE2C:hygMX at Locus2; ΔREIII::REIII(Δs1, Δs2), *swi6E(74-80)A*:KANMX | Fig. 3I; SFig5D | This Study |
| PAS 1211 (isolate 1, 2, 3) | h90, cenH:: ade6p:SF-GFP (Kint2); mat3m(EcoRV):: ade6p:mKO2; ade6p:3xE2C:hygMX at Locus2; ΔREIII::REIII(Δs1, Δs2), *swi6S18/24A,E(74-80)A):*KANMX | Fig. 3J; SFig5E | This Study |

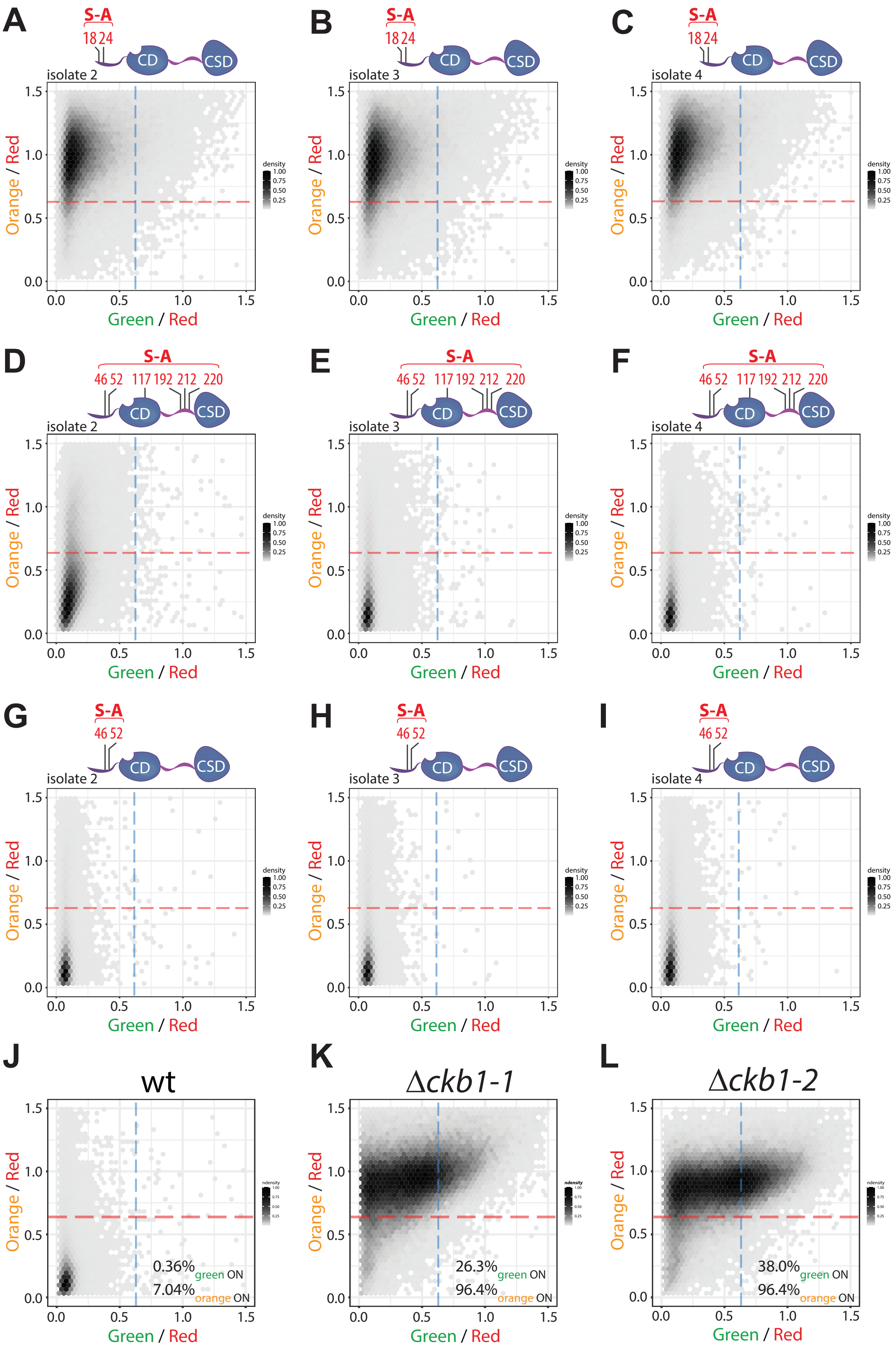


**Supporting Figure 1:** Additional isolates demonstrating that S18 and S24 in Swi6 are required for spreading, but not nucleation of heterochromatin silencing.

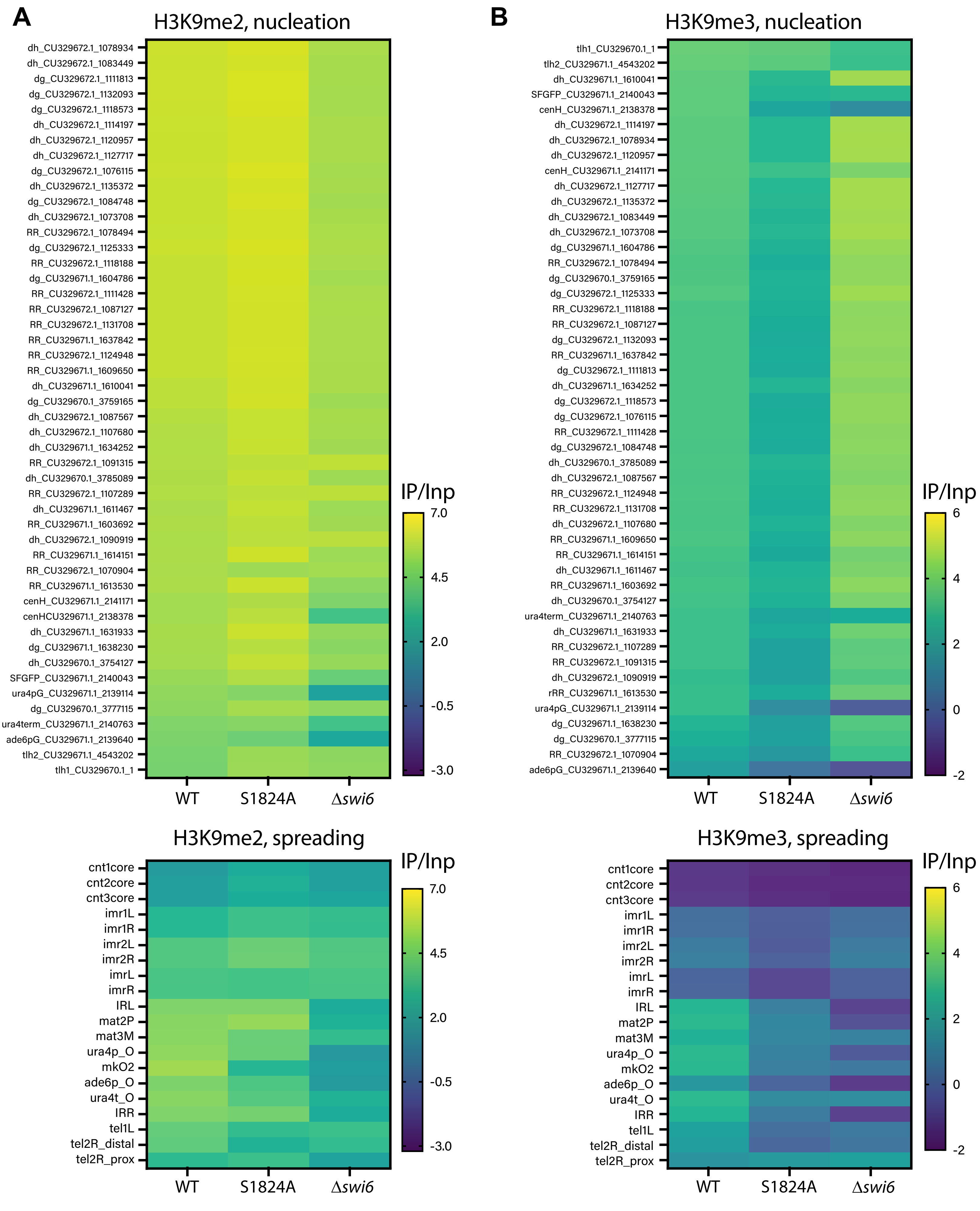


**Supporting Figure 2:** Heatmaps of H3K9me2 and H3K9me3 ChIP-seq.

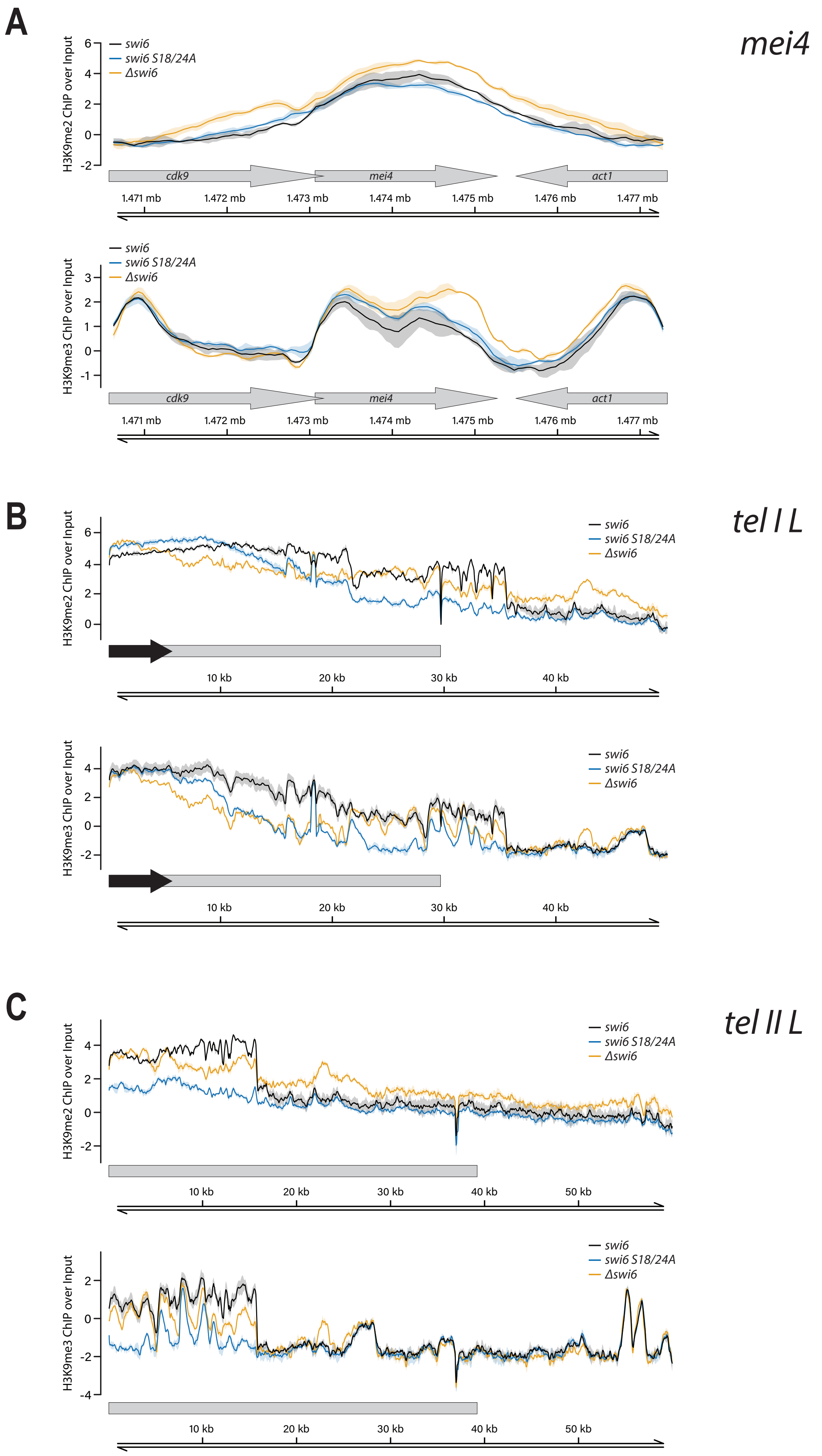


**Supporting Figure 3:** H3K9me2 and H3K9me3 ChIP-seq plots in additional genomic loci in wildtype or *swi6*$^{S18/24A}$.

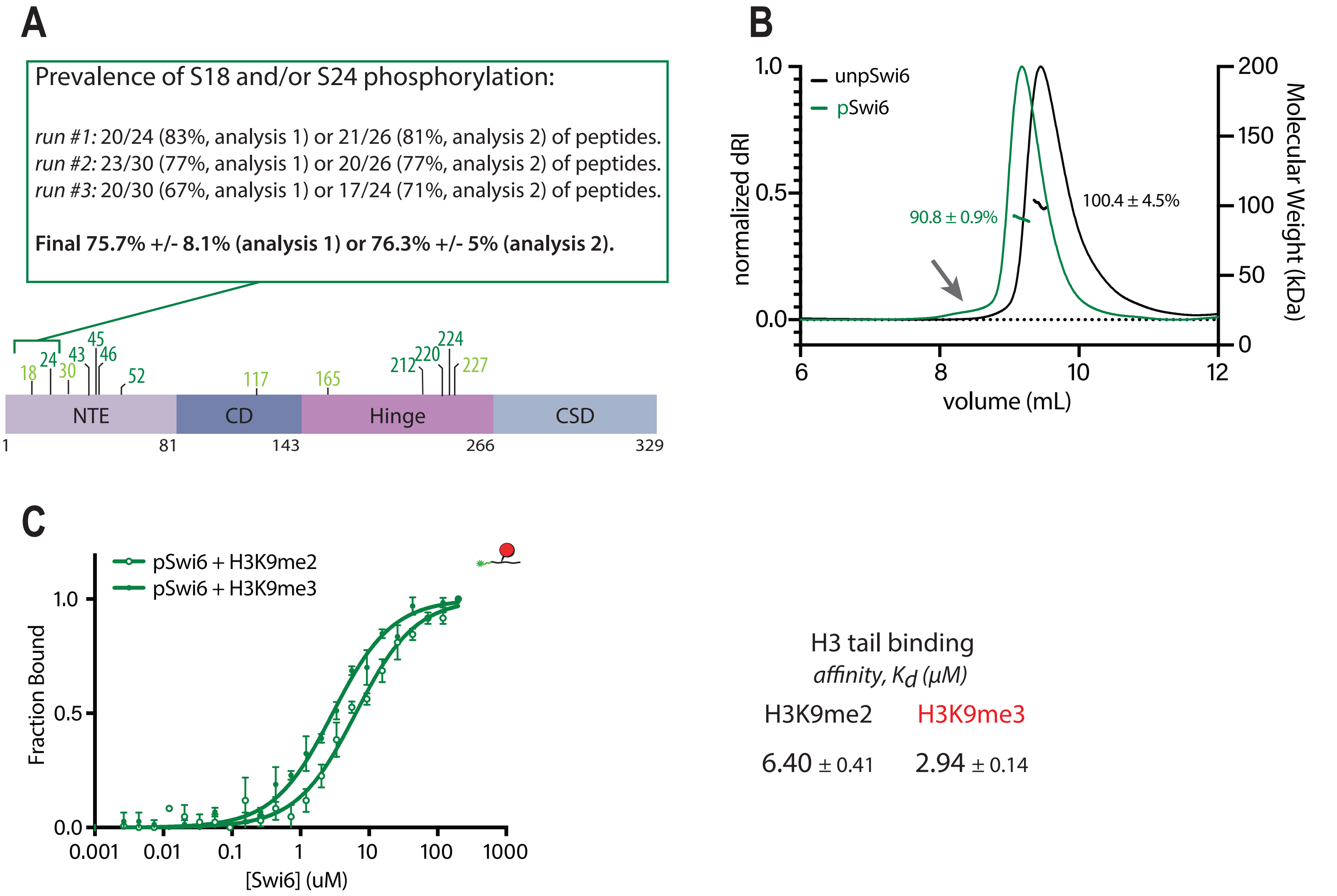


**Supporting Figure 4:** Characterization of recombinant pSwi6

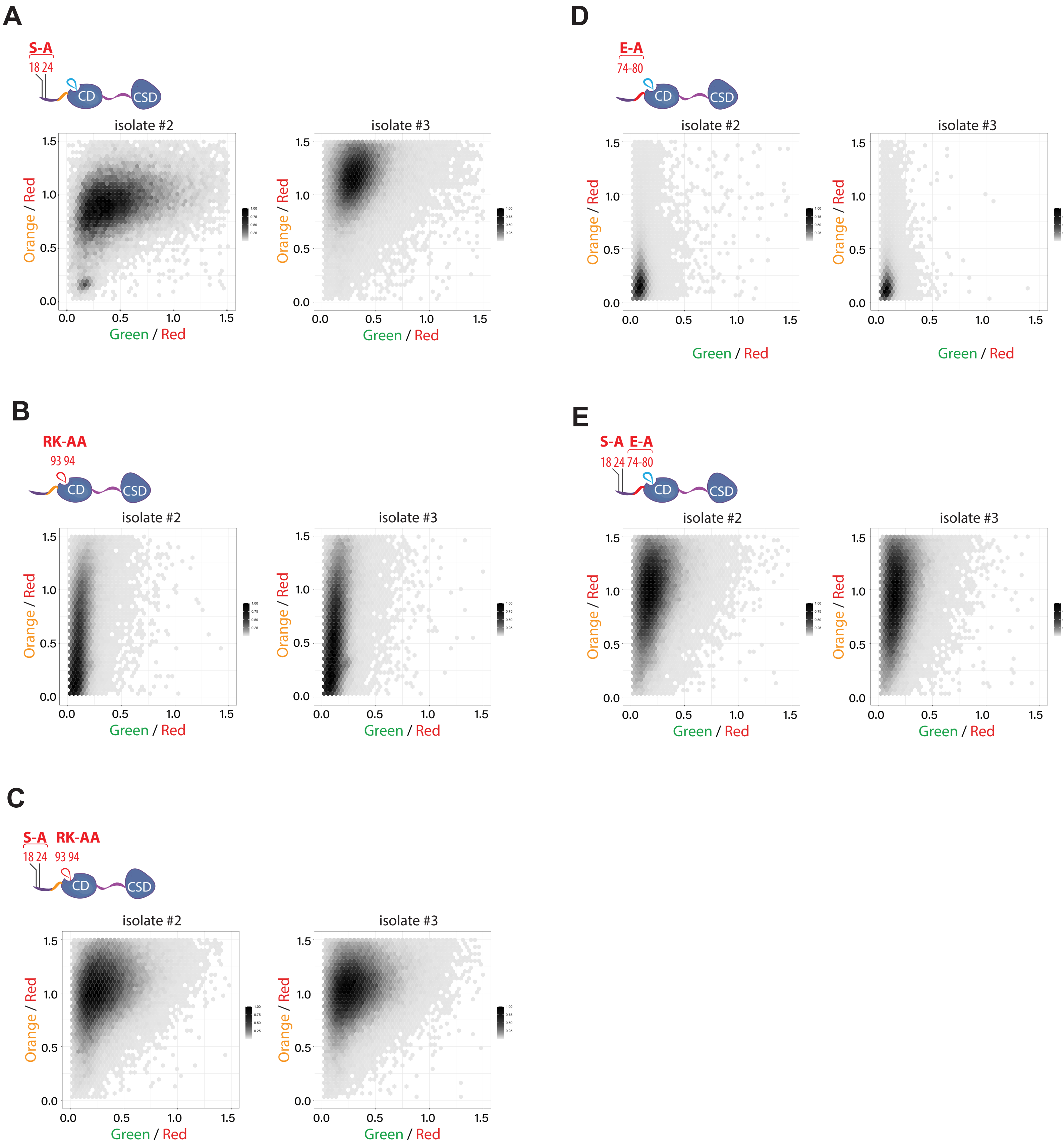


**Supporting Figure 5:** Additional isolates demonstrating that S18/S24 phosphorylation is working through, or in parallel to, known Swi6 oligomerization surfaces.

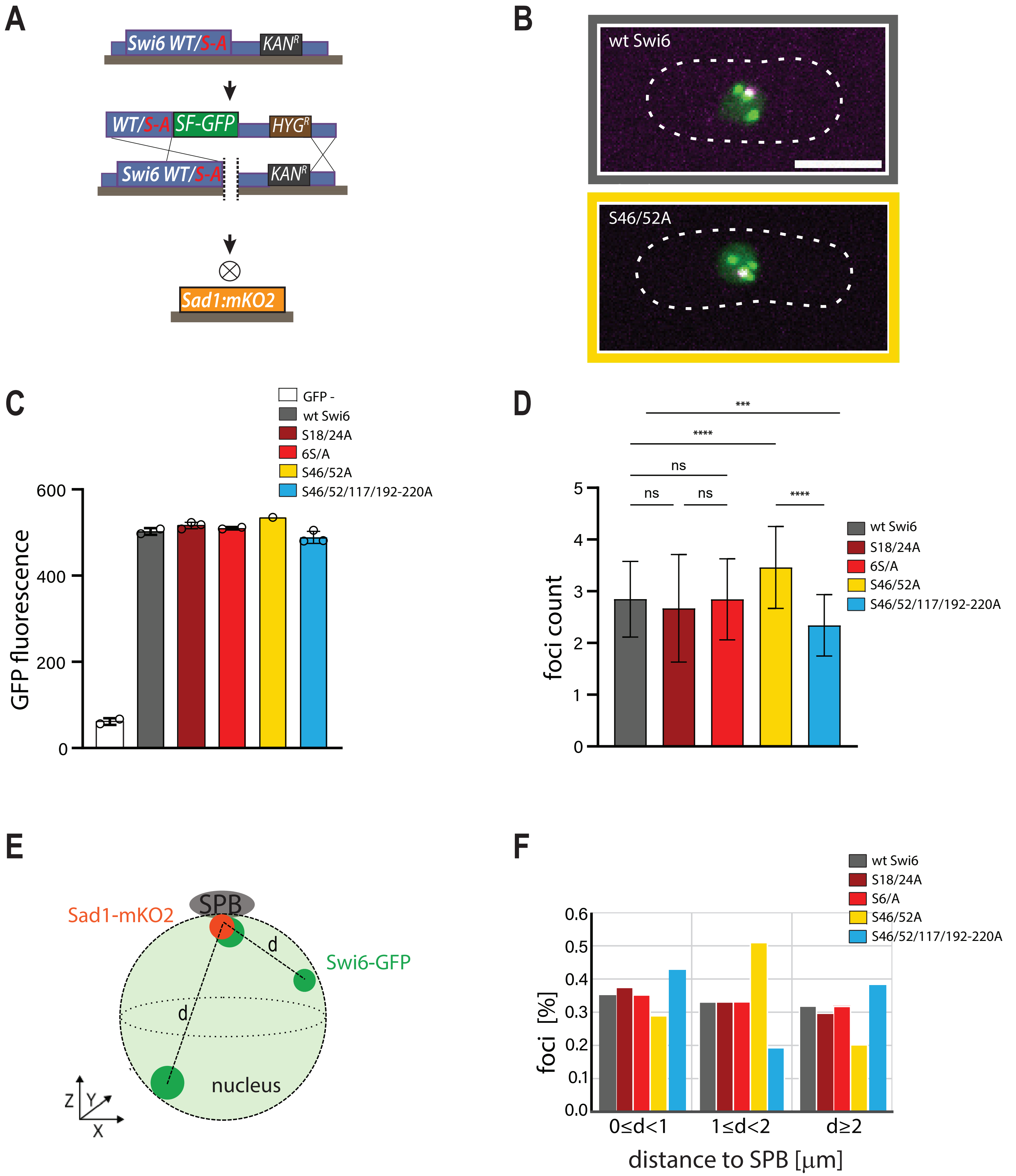


**Supporting Figure 6:** Analysis of Swi6-GFP heterochromatin foci number and spatial distribution.

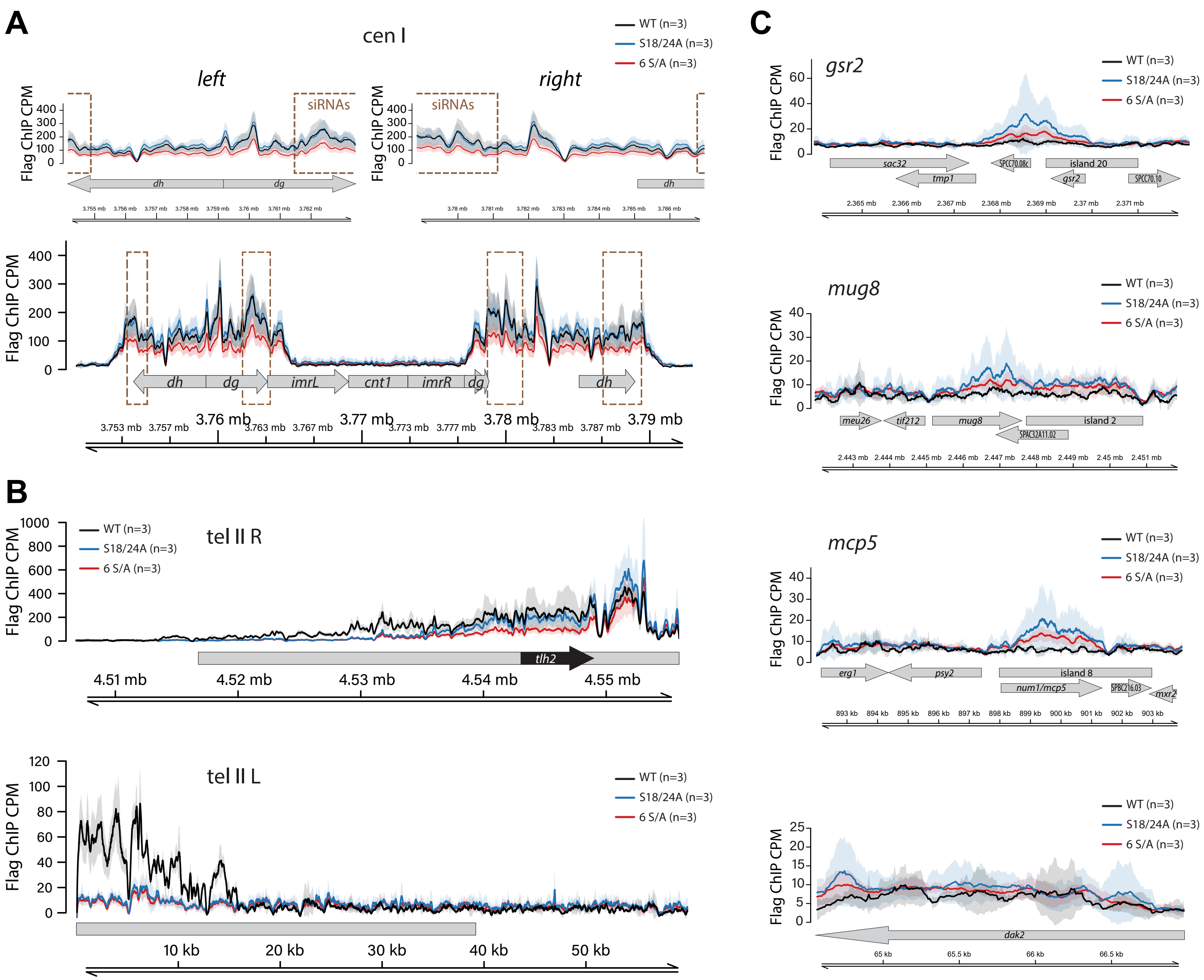


**Supporting Figure 7:** FLAG-Swi6 ChIP-Seq plots in additional genomic loci in FLAG-Swi6, FLAG-Swi6$^{S18/24A}$, and FLAG-Swi6$^{6S/A}$ strains.

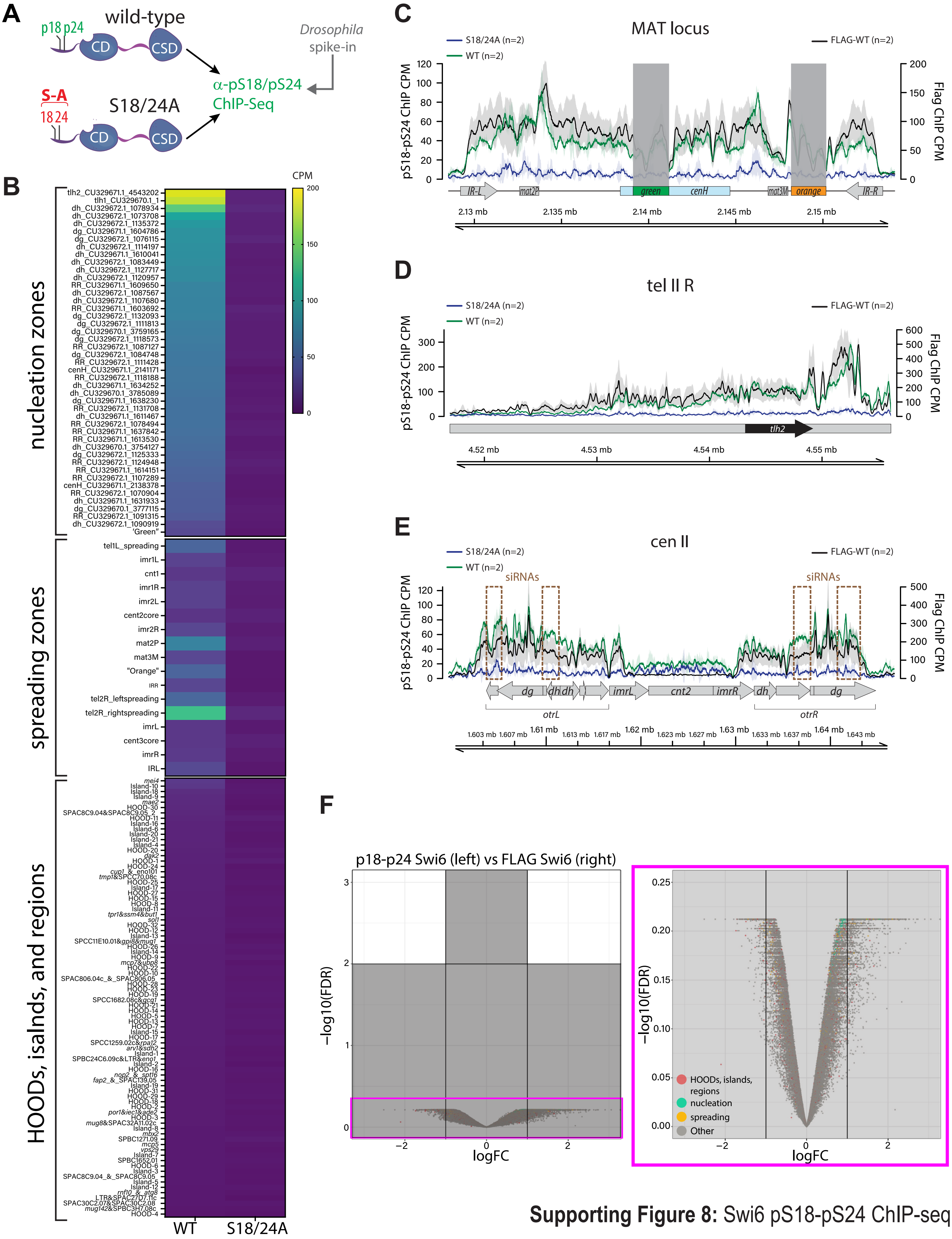


**Supporting Figure 8:** Swi6 pS18-pS24 ChIP-seq

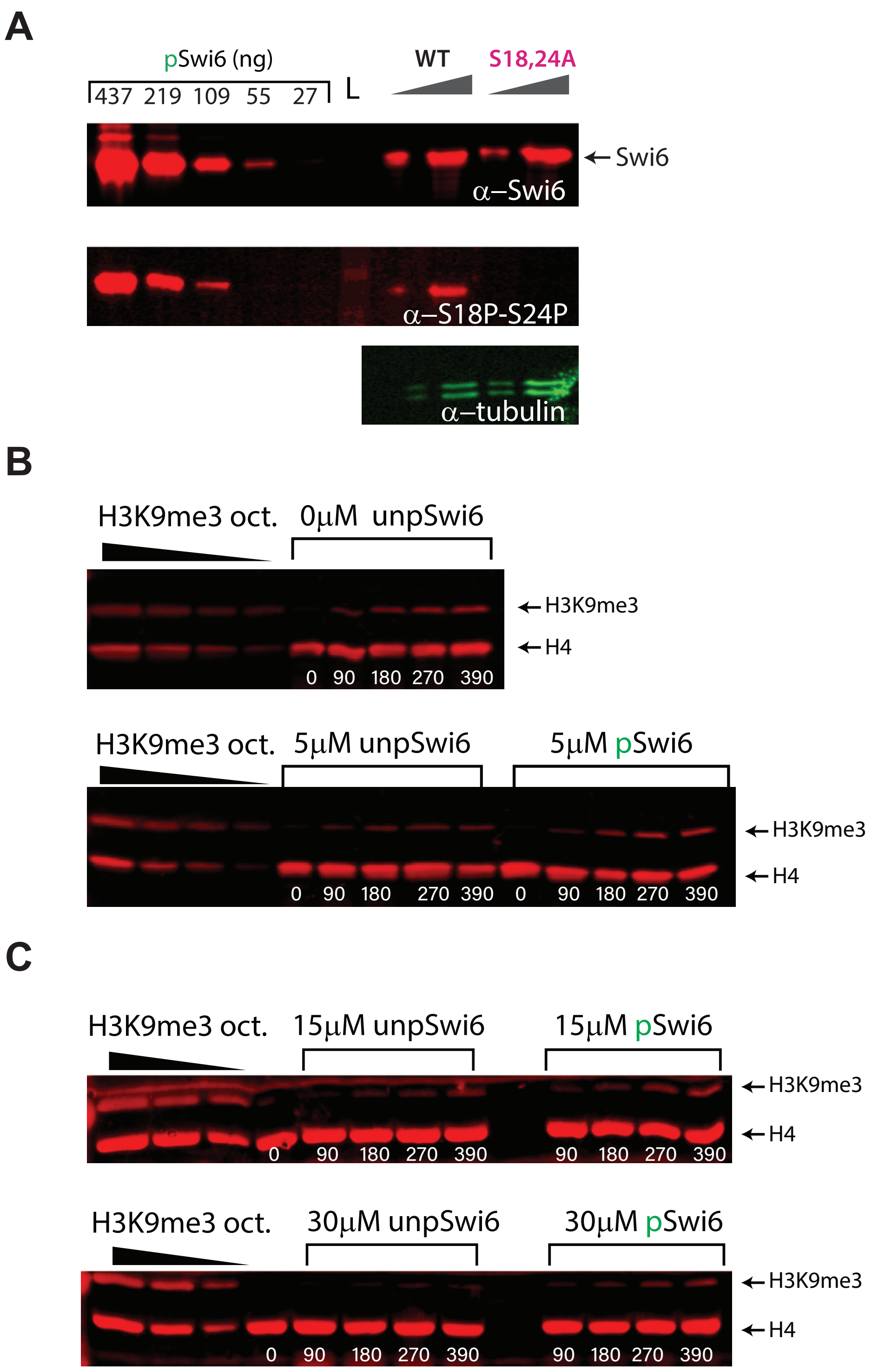


**Supporting Figure 9:** Additional replicates of Swi6 westerns from cell lysates and nucleosome trimethylation assay.

A B

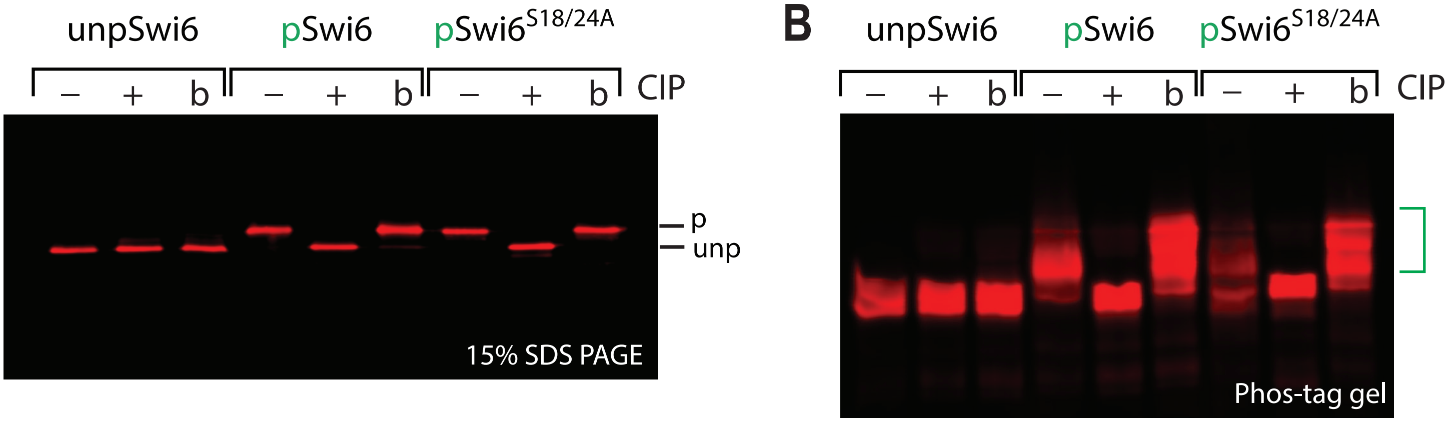


C

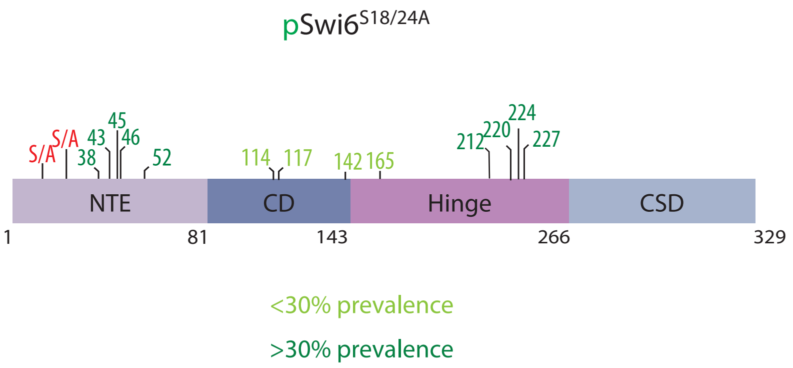


D

| | Mononucleosome binding | |
|---|---|---|
| | relative affinity | |
| | $H3K_C9me0$ | $H3K_C9me3$ |
| pSwi6 | 7.48 ± 1.07 | **1 ± 0.1** |
| pSwi6$^{S18/24A}$ | 1.67 ± 0.15 | 0.38 ± 0.003 |
| | specificity | |
| pSwi6 | 7.48 | |
| pSwi6$^{S18/24A}$ | 4.39 | |

**Supporting Figure 10:** pSwi6$^{S18/24A}$ mutant characterization.
